# Joint analysis of microsatellites and flanking sequences enlightens complex demographic history of interspecific gene flow and vicariance in rear-edge oak populations

**DOI:** 10.1101/2021.07.12.452011

**Authors:** Olivier Lepais, Abdeldjalil Aissi, Errol Véla, Yassine Beghami

## Abstract

Inference of recent population divergence requires fast evolving markers and necessitates to differentiate shared genetic variation caused by ancestral polymorphism and gene flow. Theoretical research shows that the use of compound marker systems integrating linked polymorphisms with different mutational dynamics, such as a microsatellite and its flanking sequences, can improve estimation of population structure and inference of demographic history, especially in the case of complex population dynamics. However, empirical application in natural populations has so far been limited by lack of suitable methods for data collection. A solution comes from the development of sequence-based microsatellite genotyping which we used to study molecular variation at 36 sequenced nuclear microsatellites in seven *Quecus canariensis* and four *Quercus faginea* rear-edge populations across Algeria, to decipher their taxonomic relationship, past evolutionary history and recent demographic trajectory. First, we compare the estimation of population genetics parameters and simulation-based inference of demographic history from microsatellite sequence alone, flanking sequence alone or the combination of linked microsatellite and flanking sequence variation. Second, we measure variable importance from random forest approximate Bayesian computation to identify which of these sequence types is most informative. Whereas the analysis of microsatellite variation alone indicates recent interspecific gene flow, additional information gained from combining with nucleotide variation in flanking sequences, by reducing homoplasy, suggests ancient interspecific gene flow followed by drift in isolation instead. The weight of each polymorphism in the inference also demonstrates the value of linked variations with contrasted mutation dynamic to improve estimation of both demographic and mutational parameters.

## Introduction

The build-up of species neutral genetic diversity thought time is driven by the balance between recombination, mutation and drift and its spatial distribution structured by dispersal and gene flow. As these processes are tightly linked together, understanding of mutational processes can help inform evolutionary processes better. This is especially important when studying recently diverged populations or species where the relative contribution of divergence by drift or natural selection and gene flow need to be assessed to predict species evolutionary trajectory. Recent developments of sequence-based microsatellite genotyping (Darby *et al*., 2016; De Barba *et al*., 2016; Vartia *et al*., 2016) have improved our understanding of molecular variation at microsatellite repeats and in surrounding flanking sequences (Barthe *et al*., 2012; Viruel *et al*., 2018; Lepais *et al*., 2020). Several studies demonstrated that jointly accounting for linked microsatellite and flanking sequence substitutions provides complementary information from polymorphism evolving at different rates and by different mechanisms (Hey *et al*., 2004; Ramakrishnan and Mountain, 2004; Payseur and Cutter, 2006). Assuming complete linkage due to short physical distance, linked polymorphisms share the same genealogical history. As a result, microhaplotypes combining fast evolving microsatellite repeat variations with the much slower evolving flanking sequence, should provide refined insights about past demographic history with improved estimation of temporal range and resolution (Ramakrishnan and Mountain, 2004).

For instance, Hey et al. (2004) adapted the Isolation with Migration model (IMa) to compound marker linking flanking sequence haplotype and microsatellite (HapSTR) with an illustration on two Cichlid species who recently diverged with gene flow. Using one such compound marker system improved the inference of the demographic parameters that were difficult to estimate using a single polymorphism alone. They highlighted additional avenues for progress, such as releasing constrains on the stepwise mutation model for the microsatellites and the need to use several independent loci in future applications. Conversely, Ramakrishnan & Mountain (2004) defined SNPSTR as one or more Single Nucleotide Polymorphism (SNP) linked to a microsatellite and demonstrated their use to estimate the timing of recent population divergence using both simulated and real data. They found the divergence time estimated from microsatellite variation on the background of the derived SNP allele to be more accurate and precise that the estimation based on the microsatellite variation alone. Their simulation showed that 20 SNPSTR markers provided a much better estimation that 100 microsatellites, and that the inference was robust to complex demographic scenarios involving gene flow or both population expansions and bottlenecks. In addition, divergence time between African and non-African human populations inferred from microsatellite variation alone was strongly underestimated compared with the estimation based on microsatellite variation linked to the derived SNP alleles (Ramakrishnan and Mountain, 2004). In this study, the use of a subset of the data only, i.e. the microsatellite alleles linked to the derived SNP allele aiming to reduce homoplasy (alleles identical by state but not by descent), led to much better inferences. This clearly illustrates the wealth of information that can be gained from contrasted mutation rates and mechanisms. Despite the clear demonstrated potential of using contrasted polymorphisms, these compound marker systems have been rarely studied because their genotyping is challenging (Mountain et al., 2002; Šarhanová et al., 2018; Barthe et al., 2012). This technical limitation does not hold true anymore thanks to the development of sequence-based microsatellite genotyping that greatly improves the characterisation of microsatellite and linked flanking sequences (Bradbury *et al*., 2018; Viruel *et al*., 2018; Curto *et al*., 2019; Layton *et al*., 2020). However, now that data from numerous marker systems can be easily collected at population scale, development of inferential approaches that harness this new source of information is needed (Payseur and Cutter, 2006).

Trees are a model of choice to study broad-scale postglacial colonisation pattern following the end of the last glacial maximum (Hewitt, 1999). Their unique life history traits, such as long generation time, and the use of uniparentally inherited chloroplast DNA markers, has allowed to trace glacial refugia and postglacial recolonization routes (Petit *et al*., 2002) or even Tertiary geological events (Magri *et al*., 2007) back. Nevertheless, studies conducted at a more local scale in regions known to harbour glacial refugia identified fine-scale population genetic structure within refugia (Gómez and Lunt, 2007), highlighting the value of more spatially focused studies to understand complex evolutionary processes such as vicariance and secondary contact and gene flow at play within these relict populations throughout the Pleistocene and the Holocene (Rodríguez-Sánchez *et al*., 2010; Feliner, 2014). While the Mediterranean basin is a major centre of biodiversity for the European biota, studies remain scarce on the North African side of the Mediterranean basin that harbour biodiversity hotspot and potential glacial refugia (Médail and Diadema, 2009; Lepais *et al*., 2013; Feliner, 2014). Understanding evolutionary processes in the region is particularly pressing as anthropogenic pressure is rising and increases threats posed by ongoing climate change (Hampe and Petit, 2005). *Quercus canariensis* Willd. and *Quercus faginea* Lam. are two oak species that are a good illustration of the complexity of evolutionary processes that shape present-day biodiversity within species and the threats faced by rear-edge populations. While the taxonomy of *Q. canariensis* is well defined, the taxonomic status of *Q. faginea* has been debated among taxonomists with two sub-species (*broteroi* and *faginea*) in Algeria, among the four subspecies described in *Q. faginea*, differing by morphological and environmental characteristics (Aissi *et al*., 2021). The origin of such high level of intraspecific variability is unclear and might involve a combination of genetic drift in isolated populations (Moracho *et al*., 2016) and interspecific gene flow. Localized at the rear-edge of the distribution range of the species in area of putative refugia and regional hotspot of biodiversity (Médail and Diadema, 2009), these populations are under the direct threat of contemporary environmental changes. A better understanding of the origin of the intra-specific variability and the recent demographic trajectory is paramount to evaluate the evolutionary value and conservation priority status of these populations.

The aim of this study is to assess the value of the joint analysis of microsatellite and flanking sequence variations for population genetic inference in the case of populations or species with a complex history of drift and gene flow. As a proof of concept, we studied molecular variation at 36 sequenced microsatellites across seven *Q. canariensis* and four *Q. faginea* rear-edge populations to shed light on their taxonomic status, past evolutionary history and recent demographic trajectory. Given long pollen dispersal distance, large effective population size and long generation time typical of wind-pollinated tree species such as oaks, we expect high local genetic diversity and low regional genetic structure at neutral nuclear loci. Alternatively, limited pollen dispersal between small remnant forest stands isolated by habitat fragmentation might have led to genetic drift and reduced effective population size, resulting in low local genetic diversity and high regional genetic differentiation typical of relict populations (Hampe and Petit, 2005). These two putative demographic histories can be further confounded by historical or contemporary interspecific gene flow that will additionally blurs species limits and contribute to reshuffle genetic diversity across populations. We first compared pattern of molecular variation and genetic structure observed from different polymorphisms (variation of microsatellite repeat number, SNP in flanking sequence, allele size, whole polymorphisms combined as microhaplotype) using genetic diversity and differentiation indices to illustrate the correlation or complementarity and the resolution power of different polymorphisms. We expect homoplasy within microsatellite loci to blur species limits which should be more sharply resolved when accounting for all polymorphisms.

To identify the most likely combination of historical and contemporary processes that shape current population genetic structure, we used a simulation-based inference approach that is flexible enough to model specific linked-polymorphisms observed at each locus for alternative past demographic scenarios. We derived specific summary statistics, synthetizing population-level molecular variation, that account for linked microsatellite and flanking sequence polymorphisms. We then applied a Random Forest implementation of approximate Bayesian computation (ABC) that retrieves the information content of each summary statistics and of each type of polymorphism to differentiate between concurrent demographic models and to estimate mutational and demographic parameters for the most likely demographic scenario. For historical model selection, we expect summary statistics derived from the combination of microsatellite and flanking sequence substitution to be less sensitive to homoplasy. This should help differentiate shared ancestral polymorphisms from interspecific gene flow better. For demographic parameter estimations, we expect flanking sequence SNPs to be more informative about ancient population history while fast evolving microsatellites should be more informative about recent population demography.

## Materials and methods

### Biological model and sampling

*Q. canariensis* is listed as Near Threatened at European level in the IUCN’s Red List (García Murillo and Harvey-Brown, 2017) due to population decline and increasing drought within the Iberian Peninsula. Globally, the species is considered as Data Deficient because the species lacks information on population size and status across its range especially in North Africa (Gorener *et al*., 2017). The species is present in forest stands across Morocco, Algeria and Tunisia (Figure 1a). In Algeria, *Q. canariensis* is located in either vast stands near the coastal area or in a few isolated populations in more remote southern mountain locations. *Q. faginea* is listed as Least Concern both at the European scale (Harvey-Brown *et al*., 2017) and globally (Jerome and Vasquez, 2018) due to widespread and stable abundance within its main range in Spain, Portugal and to a lesser extent in Morocco (Figure 1a). In Algeria however, *Q. faginea* is only found in four locations in the form of small and isolated populations of a few hundreds of trees including scattered individuals in mountainous landscape (Aissi *et al*., 2019).

**Figure 1:**
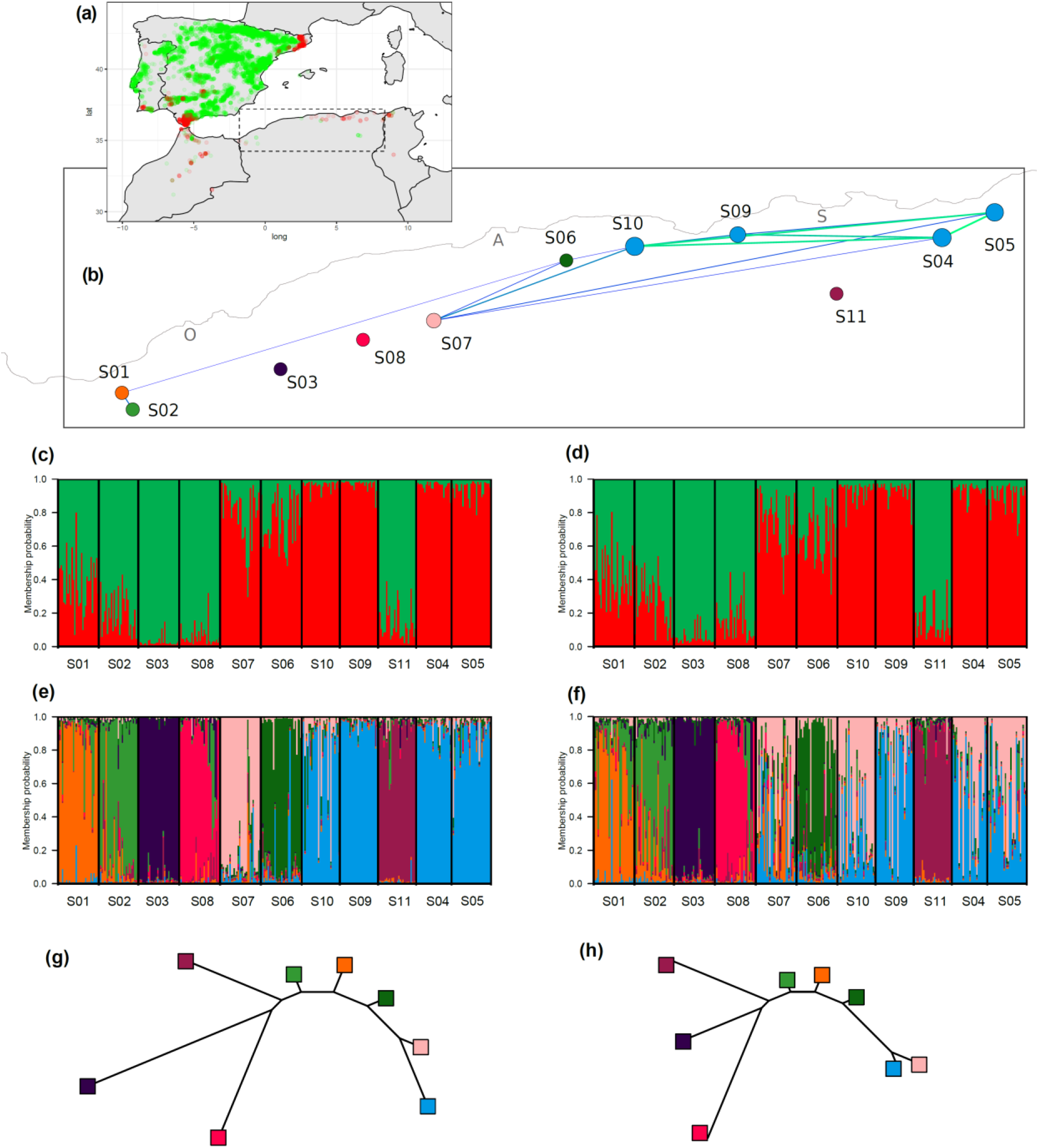
Geographical context and population genetic structure of *Quercus faginea* and *Q. canariensis* Algerian populations: (a) distribution range of *Q. faginea* (green) and *Q. canariensis* (red) estimated using GBIF occurrence data, with the studied rear-edge populations identified by the dotted rectangle; (b) EDENetwork network graph projected on a geographic map with nodes indicating spatial location of the sampled sites, node size is proportional to the number of connection with other nodes and colours reflect the dominant genetic cluster, edges link population with a pairwise genetic differentiation (Fst) below 0.04 with edge width and greenness inversely proportional to genetic differentiation. O: Oran, A: Alger and S: Skikda; (c-h) Bayesian genetic clustering Structure results using whole haplotype information (c,e,g) or the number of repeats at microsatellite only (d,f,h), assuming two (c,d) or eight (e,f) genetic clusters with the phylogenetic tree (g,h) illustrating the allele frequency divergence (Structure’s Net nucleotide distance) among the eight genetic clusters.

Fresh leaves from thirty individuals were sampled in seven *Q. canariensis* and four *Q. faginea* populations across Algeria (Supporting Material File 1 Table S1; Figure 1b). Geographical coordinates of each sampled individual were recorded and leaves were stored dried in silica gel until DNA extraction.

### Microsatellite genotyping by sequencing

Total DNA for each individual was extracted from silica dried leaf material using Invisorb DNA Plant HTS 96 Kit (Invitek) and sequence-based microsatellite genotyping (SSRseq) was used to analyse polymorphism at 36 loci as described elsewhere (Lepais *et al*., 2020). In short, 60 genomic and EST-derived microsatellites (Kampfer *et al*., 1998; Durand *et al*., 2010) were co-amplified in a single multiplexed PCR and sequenced on an Illumina MiSeq sequencer (Supporting Material File 1). After sequence demultiplexing, a bioinformatics pipeline (Lepais *et al*., 2020) integrating the FDSTools analysis toolkit (Hoogenboom *et al*., 2016) was used to convert raw sequence into genotypic data (microhaplotypes) integrating all polymorphisms identified within the microsatellite itself (number of repeats, SNP and insertion-deletion (indel)) and in its flanking sequences (SNP and indel). The bioinformatics pipeline also compares genotypes from blind-repeated genotyping of 48 individuals to estimate allelic error rate and compute the overall missing data rate for each locus. These data quality metrics are necessary to optimise the bioinformatics analysis strategy with identification of loci that cannot be reliably genotyped or for which the genotyping accuracy is only acceptable when analysing the polymorphism within the repeat motif itself (i.e. not accounting for the polymorphism in the flanking sequence (Lepais *et al*., 2020)). For each locus, every allele differing from the other by any polymorphism type was coded under an arbitrary three-digit scheme with a unique number assigned to each microhaplotype per locus.

### Multilocus genotypic dataset quality control

Assuming Hardy-Weinberg equilibrium, we tested for heterozygote deficit or excess for each locus in each population using Genetix (Belkhir *et al*., 2004) to identify outlier loci presenting extreme inbreeding coefficient that could indicate presence of null allele (positive Fis) or paralogs (negative Fis).

As clonal reproduction has been reported for oaks in harsh environmental conditions (Alberto *et al*., 2010), we identified clones using Colony (Wang, 2016) and kept only one genotype from each detected genet.

The studied loci were partly derived from expressed transcripted sequences (Durand *et al*., 2010) or selected to differentiate *Q. robur* and *Q. petraea* (Scotti-Saintagne *et al*., 2004; Lepais *et al*., 2006), therefore all may not behave neutrally. Bayescan v2.1 (Foll and Gaggiotti, 2008) was used to identify loci departing from neutrality as most of the following analyses assume neutral genetic evolution. Bayescan was run at the population level within species and at the species level using the microhaplotype dataset.

### Population genetic diversity and structure

Gene diversity (Nei’s unbiased heterozygosity), allelic richness after rarefaction to 16 diploid individuals, Fis and Fst were computed for each sampled location with FSTAT (Goudet, 1995), private alleles were computed using HP-RARE (Kalinowski, 2005), genetic differentiation accounting for allele distance in number of repeat at microsatellite alleles (Rst) was computed using SPAGeDi (Hardy and Vekemans, 2002) and genetic differentiation between species tested using 10,000 permutations of individuals between species.

To estimate genetic differentiation between populations, four different genotypic datasets were considered, accounting either for the whole haplotype, the flanking sequence haplotype, the number of microsatellite repeat or the whole allele size, to highlight potential difference in genetic structure between polymorphisms. Statistical significance for difference of genetic diversity and differentiation within species estimated from different datasets was tested by comparing locus-level statistics using a Mann-Whitney test.

Structure v 2.3.4 (Pritchard *et al*., 2000; Falush *et al*., 2003) was used to explore genetic structure between and within species. We applied the Admixture model and the Correlated Allele Frequency model with 20 replicated runs for an assumed number of genetic clusters (k) ranging from 1 to 12. To remove runs that did not converged and produced outlier results, the 15 most likely runs for each k value were kept for further inspection. Structure Harverster (Earl and vonHoldt, 2012) was used to inspect likelihood trends of the runs and Clumpak (Kopelman *et al*., 2015) to identify and filter-out sub-optimal runs.

Using the most informative whole haplotype genotypic dataset, contemporary effective population size was estimated for each sampled location using the linkage disequilibrium method implemented in NeEstimator v2.1 (Do *et al*., 2013) assuming random mating, removing alleles with frequency below 0.02 (Waples and Do, 2010) and using the parametric chi-squared method to estimate 95% confidence interval.

In addition, the geographical context of the population genetic structure was illustrated using a network graph built with EDENetworks (Kivelä *et al*., 2015) with node location representing the geographical coordinates of sampled site locations and edges linking sites showing pairwise genetic differentiation values (Fst) lower than 0.04 (determined by automatic thresholding), with edge width inversely proportional to Fst estimates.

### Population demographic history

While results from the Structure analysis show that each of the 11 studied populations can be unambiguously assigned to one of the eight delineated genetic clusters (see Results), each population cannot be unambiguously assigned to one of the two species due to genetic continuity of the studied populations. We thus investigated the most-likely demographic history that led to the observed continuous pattern of genetic differentiation. One hypothesis is that the populations diverged separately from the main core populations and that different levels of genetic drift across populations led to a continuous gradient of genetic differentiation that blurs species delineation. A second hypothesis is that gene flow taking place after a divergence period (secondary contact) produced admixed genotypes that, following recombination and drift, have mingled to form a variety of intermediate populations. While interspecific gene flow following secondary contact has been found to be the rule for sympatric European white oak species (Leroy *et al*., 2017), the studied species are not sympatric because of contrasted ecological requirements. As a result, interspecific gene flow could only happen during secondary contact or through continuous levels of low interspecific gene flow over long distances. Therefore, molecular variation was simulated for a number of realistic demographic models using the coalescent implemented in fastsimcoal2 v2.6.0.3 (Excoffier *et al*., 2013). We simulated six concurrent demographic models consisting of population divergence without gene flow (model A), population divergence with a single interspecific gene flow event (model B) or with continuous interspecific gene flow (model C) after a period of divergence without gene flow (Figure 2). Three additional models include long-term continuous interspecific gene flow between core populations of the two species (model Ab, Bb and Cb, respectively, Figure 2). Gene flow between contemporary populations and unsampled core range of the other species was modelled as directional from the core to the isolated heterospecific population because we were interested in the effect of interspecific gene flow on the studied populations specifically. In addition, we allowed interspecific gene flow only for populations showing putative admixture in Structure (Figure 1c and 1g). Effective population size was allowed to increase or decrease between the time at divergence and present. Prior distribution of demographic parameters was setup wide to reflect the lack of knowledge about the history of these populations (Table 4, Supporting Material File 1) and checked for their capacity to produce realistic genotypic data by comparing observed and simulated genotyping datasets (Cornuet, Ravigné & Estoup, 2010; Leroy et al., 2017, Supporting Material File 3). Using fastsimcoal2, we simulated realistic molecular diversity that closely matches the characteristics of each of the studied loci. We focused on two kind of variability that can be simulated: variation in repeat number at the 36 microsatellites and substitutions in the flanking sequence for the 20 loci for which flanking sequence could be reliably analysed (called HapSTR loci thereafter) and assuming complete linkage between the microsatellites and their flanking sequences. This procedure accounts for 90% of the source of variations in the dataset (Lepais *et al*., 2020), leaving out additional polymorphisms (SNP in the repeated motif or indel in the flanking sequence) that are far more complex to model due to uncertainty of the mutational mechanisms and rates.

**Figure 2:**
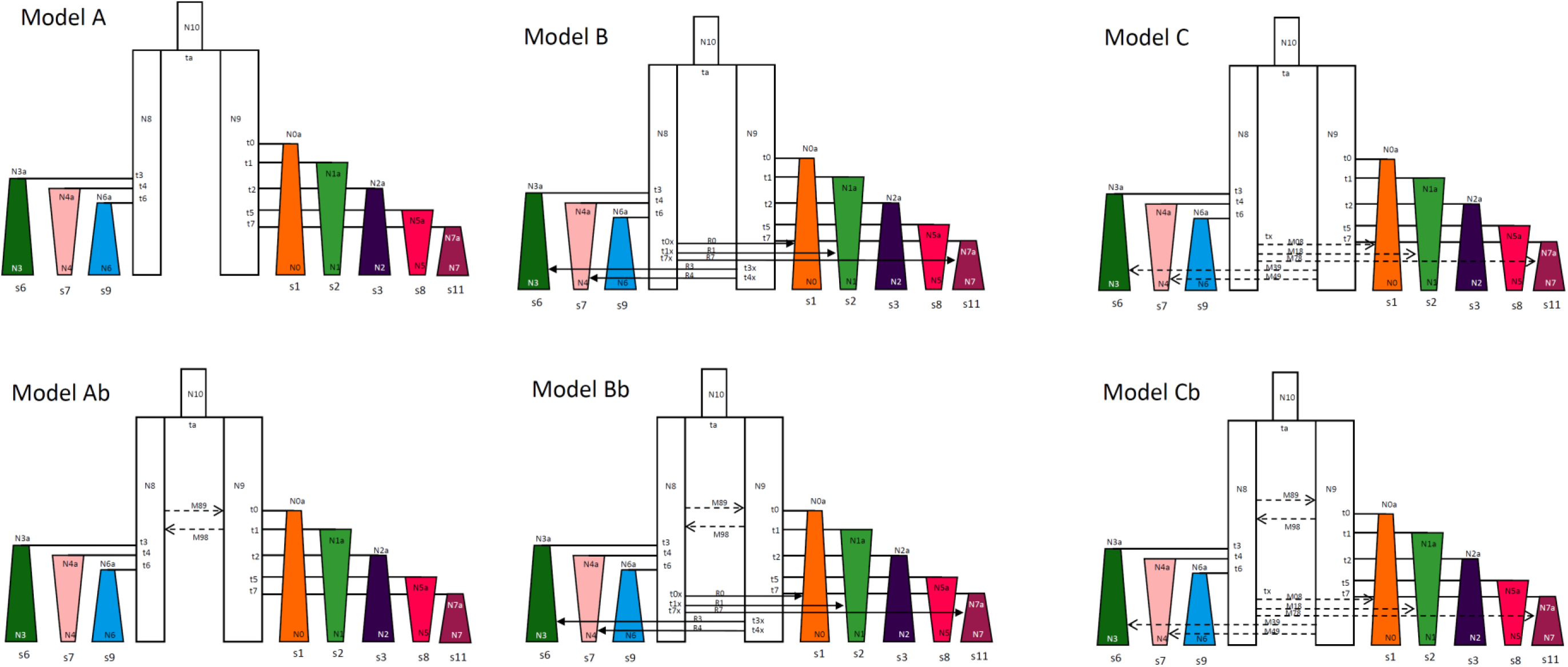
Description of simulated models of *Q. faginea* and *Q. canariensis* Algerian populations demographic history. Present-day populations diverged from their respective species main distribution area without any additional interspecific gene flow (models A and Ab), or with interspecific gene flow represented either by an instantaneous single event of interspecific gene flow (models B and Bb) or a continuous interspecific gene flow (models C and Cb). Models Ab, Bb and Cb account for long-term interspecific gene flow between species whereas models A, B and C assume that species diverged without interspecific gene flow. Unsampled (ghost and ancestral) populations are indicated in white, sampled populations are coloured according to their main Structure’s genetic cluster. Arrow with continuous line indicates single instantaneous interspecific gene flow, arrow with dashed line indicates continuous interspecific gene flow. N: effective population size, t: time of event, tx: time of secondary contact; R and M: instantaneous and continuous interspecific migration rate, s: sampled population code. Rectangle indicates fixed effective population size, isosceles trapezoid indicates variable effective population size since time of divergence; shapes and sizes are not to scale in terms of time of event or effective population size.

Each microsatellite was allowed to mutate at rates ranging from 10^−5^ to 10^−2^ mutation/generation/haploid genome (Lye *et al*., 2011) following a Generalized Mutation Model with parameter p between 0.1 and 0.5. Similarly, substitutions in the flanking sequence could mutate at rates ranging between 10^−9^ and 10^−6^ substitution/position/generation/haploid genome following neutral model of evolution. Each locus system had its own prior distributions for these parameters to model a wide range of mutational dynamics.

To compare the six alternative demographic models, 12,000 genotypic datasets were simulated for each model with demographic and mutational parameter values sampled from prior distributions. To summarize the distribution of genetic diversity among populations and locus, a total of 530 summary statistics were derived (Supporting Material File 2). We used Arlsumstat (Excoffier and Lischer, 2010) to compute 294 summary statistics that take into account the different sources of molecular variation. A first set of 98 statistics were computed with microsatellites and, when available, flanking sequences combined as microhaplotypes “considered as mutationally equidistant from each other” (standard data in Arlequin nomenclature) using the 36 loci. A second set of 135 statistics were computed based on the number of repeats from all 36 loci (microsatellite data type in Arlequin). Finally, for the 20 HapSTR loci, an additional set of 61 statistics were computed based on substitution in the flanking sequences. We then computed 224 additional summary statistics using a custom R script (Supporting Material File 4). First, 136 of these statistics were locus-level variability such as variance in allele size for the microsatellite (V), number of microsatellite alleles, number of flanking haplotypes, nucleotide diversity (theta). Second, as the covariation in diversity at linked polymorphisms might include useful information about mutational and demographic processes (Payseur and Cutter, 2006), we computed 88 statistics interrogating the association between the repeated motif number and the flanking haplotype, such as the correlation between theta and V (Payseur and Cutter, 2006) or the mean number of microsatellite alleles per flanking sequence haplotype.

We performed model choice by comparing simulation-based summary statistics from the six models to the observed summary statistics from the real dataset within the approximate Bayesian computation framework (Beaumont *et al*., 2002) and using the Random Forest approach implemented in the R package abcrf (Pudlo *et al*., 2016; Raynal *et al*., 2019).

We first built a classification random forest model using 1,000 trees and a training dataset consisting of simulation-based summary statistics obtained from all models. We estimated the classification prior error rate for each model using an “out-of-bag” procedure to estimate the power of the genetic data to differentiate between the demographic models given the model and prior specifications (Pudlo *et al*., 2016). Then, we used the summary statistics computed based on the observed genotypic data to predict the demographic model that best fit the data using a regression forest with 1,000 trees. In addition, we also repeated the model choice analysis using only the 231 summary statistics derived from the microsatellite motifs from the 36 loci.

We then used the overall most likely scenario to simulate 100,000 additional genetic datasets using parameters and prior distributions described above to estimate demographic model parameters. We built a regression random forest model implemented in the R package abcrf based on the summary statistics using 1,000 trees. The out-of-bag procedure (Raynal *et al*., 2019) was used to assess prediction error and inference power by comparing true (simulated) and estimated parameter values. We then estimated the posterior median, 0.05 and 0.95 quantiles of demographic and mutational parameters using a random forest regression model based on all summary statistics computed from the observed haplotypic dataset.

Finally, we assessed the value of different or combined polymorphisms for demographic inference by quantifying summary statistics importance for both model choice and parameter estimation (Estoup *et al*., 2018). To better interpret the significance of variable importance, we introduced 12 additional random summary statistics (“noise variable”) in the datasets (Chapuis *et al*., 2020). These consisted in four random floating point variables sampled between 0 and 1, four random floating variables sampled between 0 and 10 and four random natural number variable sampled between 1 and 30. As random variables, these noise variables should not contribute to the inference, and thus will provide a benchmark to assess the significance of the variable importance metrics reported for each summary statistics (Chapuis *et al*., 2020). Thus, summary statistics with variable importance higher than random summary statistics were considered informative.

## Results

### Genotyping

After filtering for missing data, genotyping error, Hardy-Weinberg equilibrium, neutrality and clonality (see Supporting Material File 2), the final genotypic dataset consisted in 36 high quality sequenced loci genotyped in 318 individuals. This includes 20 linked flanking sequence and microsatellite loci, called HapSTR thereafter, and 16 loci that could not be reliably analysed across the whole haplotype and for which only the microsatellite motif was analysed, called STR thereafter (Supporting Material File 2).

### Molecular variation over loci

In addition to variability for the number of microsatellite repeats, SNP and indel within the microsatellite motif and the flanking sequences was the rule rather than the exception (Supporting Material File 1 Table S2 and File 2). A total of 438 alleles differing in size would have been observed using traditional capillary-electrophoresis, which represent half of the number of alleles when considering all source of molecular variation identified using sequence data (899 haplotypes in total, Supporting Material File 2). As a result, size homoplasy (allele having identical length but differing in sequence) amounts to 51% overall (56% for HapSTR and 31% for STR). Across the 36 loci, 32 showed SNP within the microsatellite motif (for a total of 115 SNP), nine show indel within the microsatellite motif (for a total of 10 indel) and nine microsatellites consisted of more than one repeated motif (compound microsatellites). SNP among the flanking sequence for the 20 HapSTR was widespread with an average of 7.5 SNP per locus (min: 3, max: 14) and an average of one SNP every 6.7 nucleotides (15% of flanking sequence position showed a SNP). Half of the HapSTR showed an indel with an average of 0.9 indel per loci (min: 0, max: 4).

### Molecular variation across populations

Interspecific genetic differentiation (Fst) varies from 0.030 when computed based on flanking sequence haplotype to 0.045 when computed from the whole haplotype (Table 1). Differentiation among *Q. canariensis* populations (average: 0.026) was lower than among *Q. faginea* populations (average: 0.043) irrespectively of the polymorphism considered. The comparison of Fst computed from the whole haplotypes among *Q. canariensis* populations (Fst=0.026) and among *Q. faginea* populations (Fst=0.049) was statistically significant (Mann-Whitney z-score = −3.73, p-value = 0.0002, N=36). The higher differentiation among *Q. faginea* populations is mostly driven by differentiation at microsatellite variation which translated into higher Fst estimates based on allele size, repeat number or whole haplotype (Table 1). By contrast, differentiation at flanking sequence is not statistically different among *Q. canariensis* and *Q. faginea* populations (z-score = −0.35, p.value = 0.7363, N=20) and differentiation in *Q. canariensis* is similar at the microsatellite and the flanking sequence (0.027 and 0.024 respectively, Table 1).

**Table 1:**
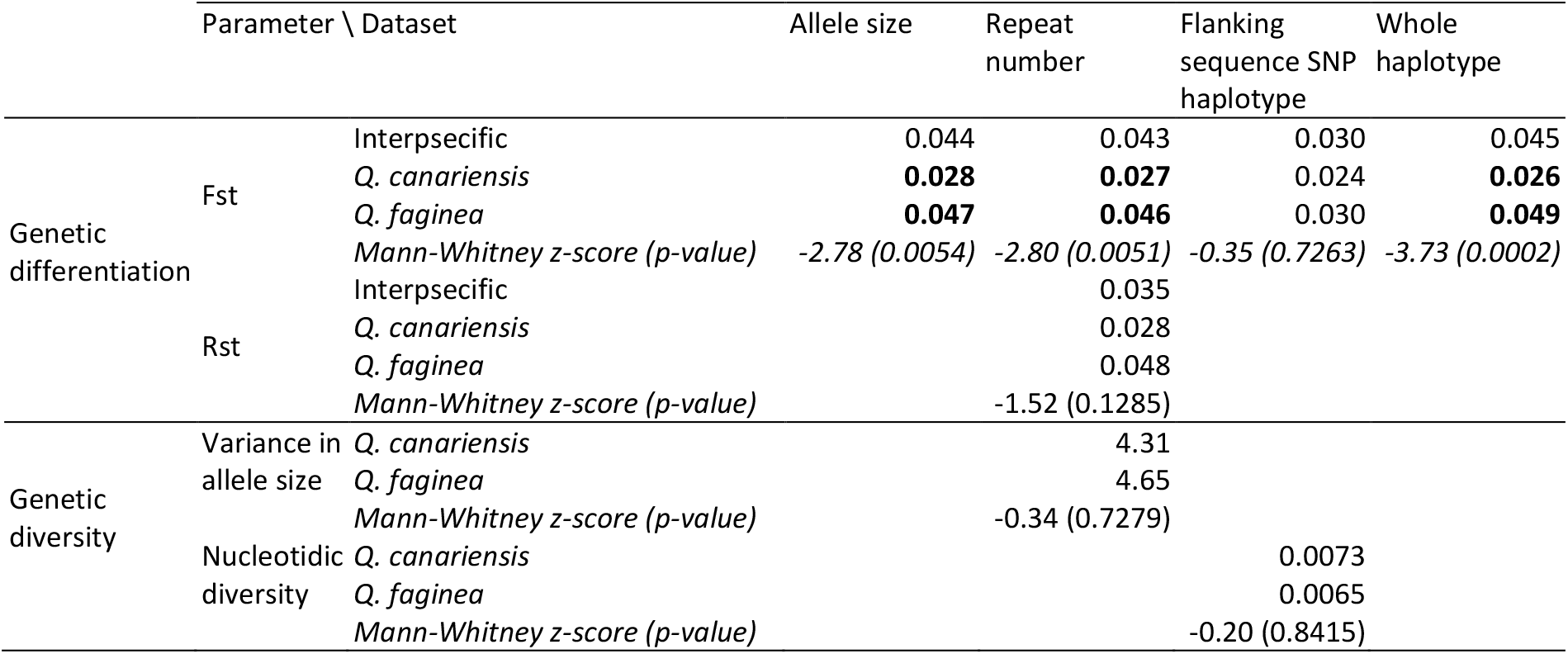
Between and within species genetic diversity and differentiation parameters computed using different polymorphism types.

Genetic differentiation based on microsatellite allele distances (Rst) was not statistically significant from genetic differentiation considering only allele identity (Fst) at the intraspecific and interspecific level (all permutation tests not significant).

Genetic diversity did not differ between species (Table 1): variance of allele size at microsatellites amount to 4.31 and 4.65 and nucleotide diversity at flanking sequence to 7.3.10^−3^ and 6.5.10^−3^ for *Q. canariensis* and *Q. faginea*, respectively. Locus-level comparison showed low and not significant correlation between variance of allele size at microsatellite and nucleotide diversity at linked flanking sequence (Pearson product moment correlation: 0.15 (t = 0.62, df = 16, p-value = 0.54) and 0.20 (t = 0.84, df = 17, p-value = 0.41) for *Q. canariensis* and *Q. faginea* respectively).

At population level, allelic richness, gene diversity and private alleles showed similar variation between populations (Figure 3), except for populations S02 and S03 that harbour higher and lower diversity, respectively (Figure 3a, 3c and 3e). Flanking sequence haplotypes consistently showed lower allelic richness and genetic diversity than STR and their combination under HapSTR showed substantially more variability (Figure 3a and 3c). This observation is explained by the lack of correlation between allelic richness or gene diversity between flanking sequence and linked STR (Figure 3b and 3d). The mean number of private alleles per locus is generally low and similar between flanking sequence and STR (0.25 and 0.20 respectively), at the exception of S02, S06 and SS09 populations were private flanking sequence were more frequent (Figure 3e). However, the combination of the two sources of variations in HapSTR showed a much higher mean number of private alleles (0.88). Overall, these observations indicate that flanking sequence and linked STR harbour complementary information.

**Figure 3:**
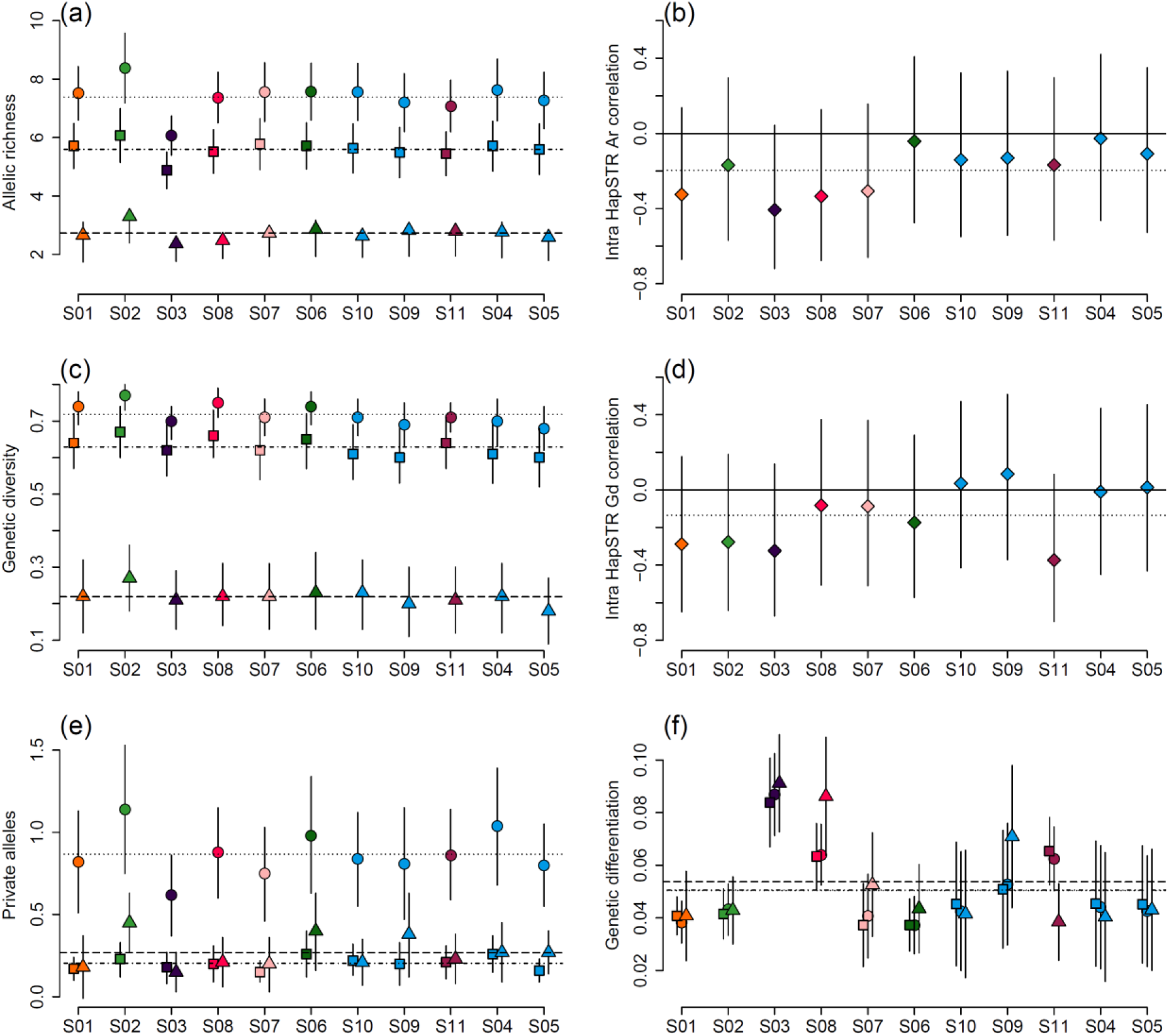
Genetic diversity of *Q. faginea* (S02, S03, S08, S11) and *Q. canariensis* (S01, S04, S05, S06, S07, S09, S10) Algerian populations: (a) allelic richness, (b) within-loci correlation between allelic richness (Ar) at individual microsatellite and their flanking sequence, (c) gene diversity, (d) within-loci correlation between gene diversity (Gd) at individual microsatellite and their flanking sequence, (e) private alleles, (f) genetic differentiation (Fst, summarized by averaging all pairwise genetic differentiation estimates that involved the focal population). Symbols represents mean over loci (square: microsatellite variation, triangle: flanking sequence variation, circle and diamond: whole haplotype integrating both kind of polymorphisms) and the bar shows estimate variability (plus and minus two standard deviations over loci). Populations are ordered from west to east along the x-axis, colours refer to Structure results, horizontal dashed lines represent the average value across populations.

Population-specific genetic differentiation is the highest for S03, S08 and S11 populations and shows the highest variability for *Q. canariensis* S04, S05, S09 and S10 populations (Figure 3f), which is consistent with *Q. canariensis* populations showing low divergence from each other and *Q. faginea* populations showing higher distinctiveness in the genetic differentiation network (Figure 3b). Genetic differentiation has similar values for flanking sequence haplotypes, STR and combined information, at the exception of a few populations where flanking sequence haplotypes showed higher (S08, S07, S09) or lower (S11) genetic differentiation (Figure 3f).

Structure analysis assuming two genetic clusters (K=2), broadly delineate the two species using either HapSTR haplotypes or STR alleles (Figure 1c et 1d respectively): *Q. canariensis* S04, S05, S09, S10 populations in the one hand and *Q. faginea* S03 and S08 populations in the other hand clearly fall in their own specific cluster. However, *Q. canariensis* S01, S06 and S07 populations and *Q. faginea* S02 and S11 populations showed admixture or unsorted ancient ancestry. While S01 had been affected as *Q. canariensis* based on morphology, it was found to be more genetically similar to *Q. faginea*. Assuming a higher number of genetic clusters resulted in a more detailed view of the intraspecific genetic structure (Supporting Material File 1 Figure S2). At K=8, using HapSTR haplotypes, results in a clear delineation of 7 populations into specific clusters in addition to a genetic cluster regrouping 4 *Q. canariensis* populations (Figure 1e). Population delineation is less clear using STR alleles, especially for *Q. canariensis* populations (Figure 1f, Supporting Material File 1 Figure S3). Differentiation between genetic clusters do not showed a bipartite hierarchical structure expected when analysing two species (Figure 1g et 1h). Instead, five of the clusters showed gradual level of differentiation starting from the *Q. canariensis* side (right part of the tree in Figure 1g), while three genetic clusters on the *Q. faginea* side showed high genetic divergence (left branch of the tree in Figure 1g). Populations with intermediate ancestry coefficient assuming two genetic clusters (S02, S01, S06, S07 and S11) now constitute independent clusters that located at intermediate positions between the two species extremities (Figure 1e). The tree topology is similar when using only STR alleles, but branches are shorter (Figure 1h).

Contemporary effective population size shows high variability with estimates ranging as low as 50 individuals for S03 and S08 population to 350 individuals for S05 (Supporting Material File 1 Figure S1). Excluding the S04 population for which sampled size is too low for a point estimate (indicating a higher effective population size with the 95% lower bound confidence interval estimated at 524 individuals), the contemporary effective population size averaged to 150 individuals over populations.

### Demographic inferences

#### Model choice

When using the information from both the microsatellite and the linked flanking sequence (HapSTR), the most likely model to explain the observed data was model Ab (posterior probability: 0.47, Table 2a) which consisted in species divergence with gene flow followed by independent divergence of populations without additional more recent interspecific gene flow (Figure 2). The out-of-bag procedure estimated that, given the choice of the model Ab and the uncertainty in differentiating the models given the data and the prior distribution settings, there is a 49% probability that the true model is indeed the model Ab, and a 18% or 17% probability that the true model is model Cb and Bb respectively (Table 2b).

**Table 2:** Random forest ABC Model choice (a) by using jointly microsatellite and flanking sequence information (HapSTR) or information derived only from the microsatellite variations (SSR only). Power to differentiate alternative demographic models for HapSTR dataset (b) given the model and prior range specifications. The table reports how many of the 12,000 simulated datasets generated under a specific scenario (rows) are classified into each demographic scenario (columns). The overall scenario classification error is computed based on the number of incorrectly classified dataset; the last column shows the percentage of simulated datasets classified as Ab, the most likely scenario for the observed genotypic dataset. Bold numbers indicate >10% incorrect classification, underlined number indicates correct classifications.

| (a) |  |  |  |  |  |  |  |  |  |
| --- | --- | --- | --- | --- | --- | --- | --- | --- | --- |
|  | Votes<br>Model A | Votes<br>Model B | Votes<br>Model C | Votes<br>Model Ab | Votes<br>Model Bb | Votes<br>Model Cb | Most<br>likely<br>model | Posterior<br>probability |  |
| HapSTR | 163 | 103 | 114 | 271 | 191 | 158 | Ab | 0.47 |  |
| SSR only | 31 | 113 | 82 | 169 | 286 | 319 | Cb | 0.40 |  |
| (b) |  |  |  |  |  |  |  |  |  |
| HapSTR-Classified<br>models | Model A | Model B | Model C | Model Ab | Model Bb | Model Cb | total | Scenario<br>classification<br>error | Probability<br>given the<br>selected best<br>model is Ab |
| HapSTR-Simulated<br>models |  |  |  |  |  |  |  |  |  |
| Model A | 10142 | 183 | 290 | 1042 | 183 | 160 | 12000 | 15.5% | 8% |
| Model B | 54 | 8017 | 2087 | 623 | 765 | 454 | 12000 | 33.2% | 5% |
| Model C | 479 | 3905 | 5905 | 562 | 520 | 629 | 12000 | 50.8% | 4% |
| Model Ab | 450 | 228 | 227 | 6685 | 2419 | 1991 | 12000 | 44.3% | 49% |
| Model Bb | 35 | 559 | 305 | 2248 | 5488 | 3365 | 12000 | 54.3% | 17% |
| Model Cb | 37 | 366 | 522 | 2380 | 3769 | 4926 | 12000 | 59.0% | 18% |
| total | 11197 | 13258 | 9336 | 13540 | 13144 | 11525 |  | 42.8% |  |

A total of 98 summary statistics had variable importance higher than randomly generated statistics, and thus were considered informative to differentiate alternative scenario (Figure 4a). These informative summary statistics represented 44% of the total variable importance. Among them, 24% derived from linear discriminant axes, 35% from summary statistics computed from microsatellite variation, 33% from haplotypes and only 8% from the substitutions within the flanking sequences (Figure 4a).

**Figure 4:**
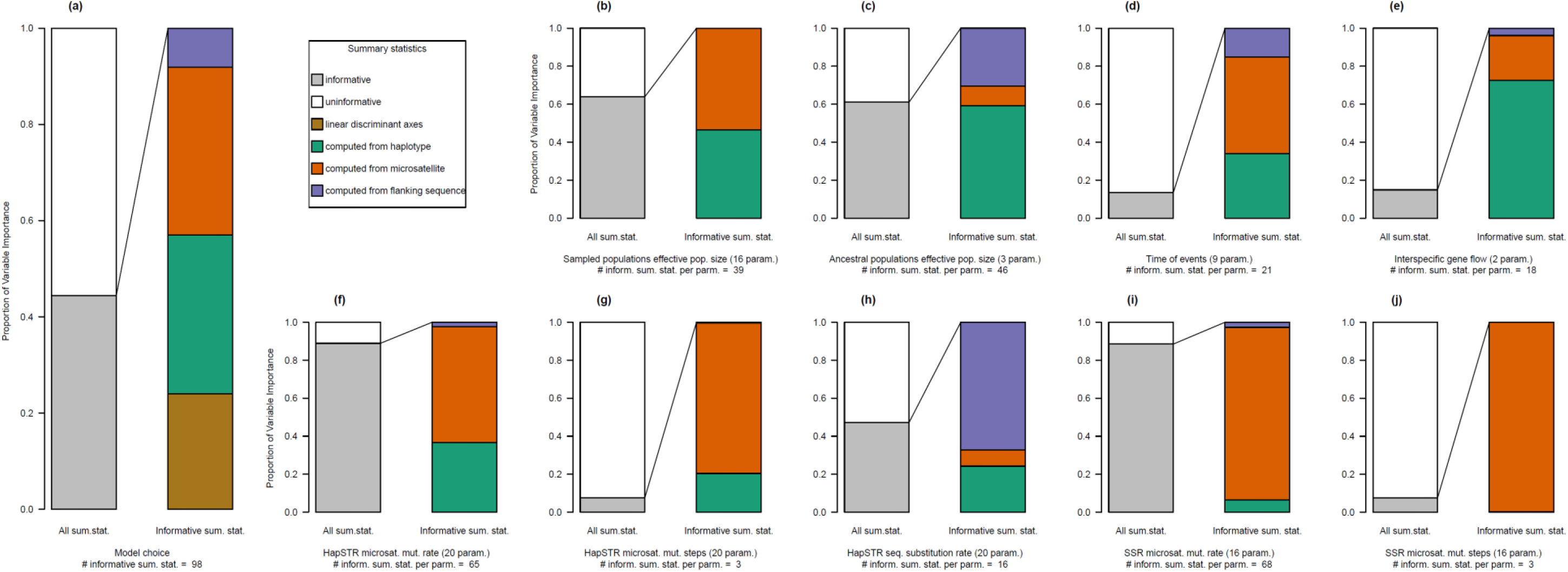
Contribution of different types of polymorphism measured as Variable Importance (y-axis) for (a) model choice and parameter estimation: (b) sampled and (c) unsampled population effective population size, (d) time of event, (e) interspecific gene flow, HapSTR microsatellite (f) mutation rate, (g) mutation steps and substitution rate (h), SSR microsatellite mutation rate (i) and mutation steps (j). In each subfigure, the leftmost bar shows the proportion of total Variable Importance resulting from informative summary statistics, while the rightmost bar shows the contribution of the different groups of summary statistics derived from different types of polymorphisms.

It should be noted that when considering microsatellite variation alone, the most likely scenario was model Cb (posterior probability: 0.40, Table 2b) which consisted in species divergence with gene flow followed by independent divergence of populations followed by interspecific gene flow after a period of strict isolation (Figure 2). However, since HapSTR integrates more molecular information, this strategy provides higher resolution (posterior probability of 0.48 and 0.40 for HapSTR and STR respectively) and power (overall classification error of 42.7% and 44.4% for HapSTR and STR respectively). Therefore, we selected scenario Ab to estimate its demographic parameters using the HapSTR dataset.

#### Demographic and mutational parameter estimations

Microsatellite mutation and substitution rates were accurately estimated (all posterior MNSE lower than 0.11; Supporting Material File 2) with an average of 65 and 68 informative summary statistics representing 89% and 88% of the total variable importance for microsatellite mutation rates at HapSTR loci and SSR loci respectively (Figure 4f and 4i) and an average of 16 informative parameters for substitution rate that represents 47% of the total variable importance (Figure 4h). On the contrary, the parameter p controlling the number of mutational steps at microsatellite was difficult to estimate (posterior MNSE between 2.6 and 7.6; Supporting Material File 2) and had only an average of 3 informative summary statistics per parameters that represented 7.6% of the total variable importance for all loci (Figure 4g and 4j).

Estimated microsatellite mutation rates varied from 2.2 × 10^−5^ to 2.1 × 10^−3^ mutation per generation per haploid genome (mean: 3.1 × 10^−4^, 95% CI: 4.5 × 10^−5^ – 2.4 × 10^−3^) and flanking sequence substitution rates varied from 4.5 × 10^−8^ to 4.8 10^−7^ (mean: 2.5 × 10^−7^, 95% CI: 2.8 × 10^−8^ – 9.0 × 10^−7^, Supporting Material File 2).

Posterior effective population size parameters showed restricted ranges compared to prior, as well as reduced estimation error (posterior NMAE below 0.32, Table 3). Contemporary and ancestral effective population sizes of sampled populations could be estimated, with an average of 39 informative summary statistics (Figure 4b) representing 64% of the total variable importance. Effective population size of unsampled populations (i.e. *Q. canariensis* and *Q. faginea* core range, and their ancestor) could be estimated with an average of 46 informative summary statistics (Figure 4c) representing 61% of the total variable importance.

**Table 3:**
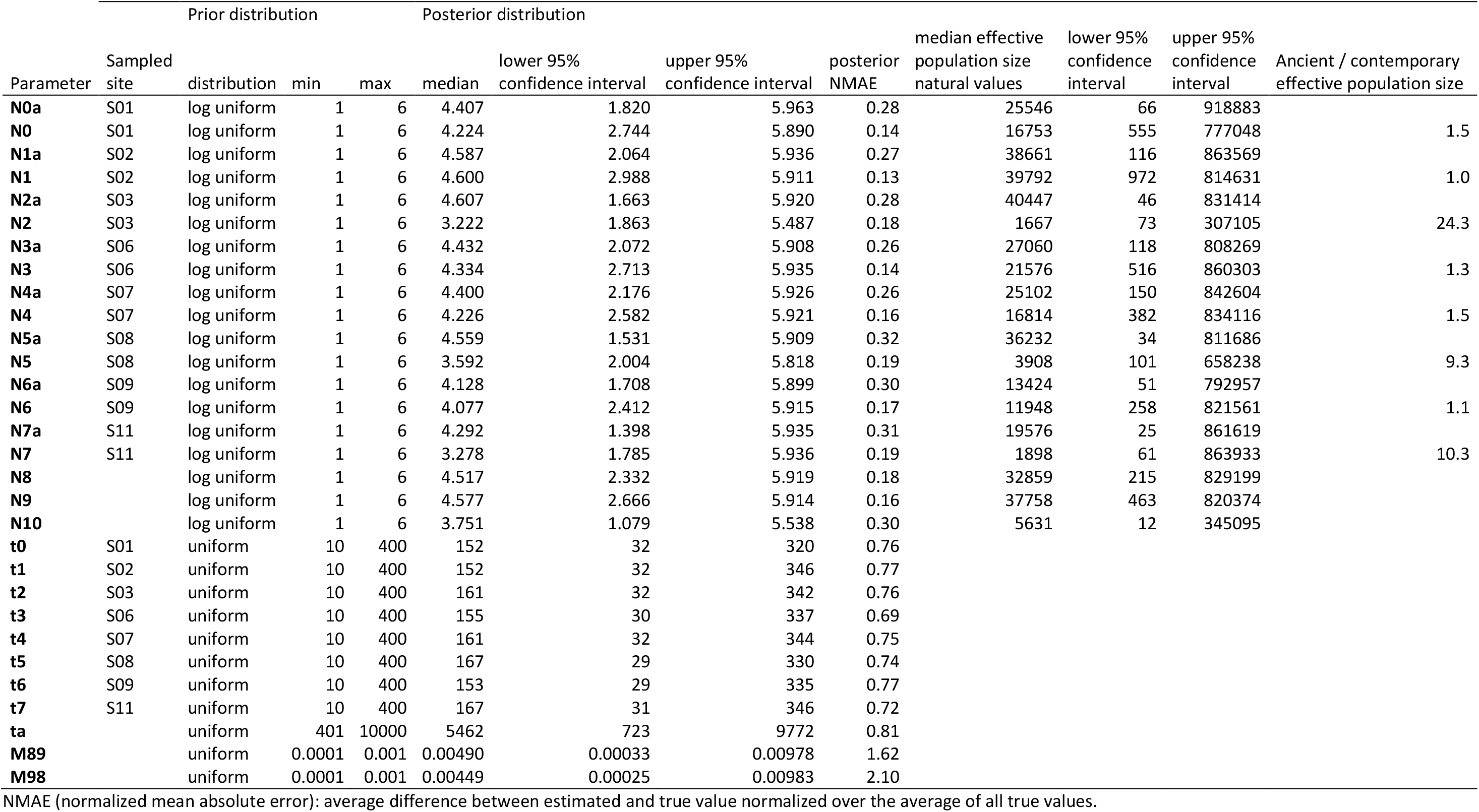
Demographic parameter prior specification and posterior estimations for the most likely model Ab.

Effective population size estimates showed some variation between populations (Table 3). Effective population size at divergence time were generally higher than recent effective population size, with clear sign of effective population size reduction for S08 (ratio N5a/N5 = 9.3, N5 = 3908 haploid genomes), S11 (ratio of N7a/N7 = 10.3, N7 = 1898 haploid genomes) and S3 (ratio of N2a/N2 = 24.3, N2 = 1667 haploid genomes). Populations that did not experience significant effective population size change had relatively moderate (~10,000 for S09, ~20,000 for S01, S06 and S07) or high (~40,000 for S02) effective population sizes.

Overall, only 36 out of the 518 summary statistics were not informative for the estimation of demographic or mutational parameters. Most of them were associated with flanking sequence diversity or divergence and correlation between diversity at microsatellite and flanking sequence (Supporting Material File 2).

## Discussion

### The complex demographic history of rear-edge populations

A first aim to this study was to assess how the informative content of microsatellite sequence can help us understand better the role of interspecific gene flow and recent demographic dynamics in structuring genetic diversity in complex population dynamics of *Q. faginea* and *Q. canariensis* at the rear-edge of their distribution as a case study. A genetic clustering analysis assuming two genetic clusters showed highly admixed populations, which can be interpreted as evidence for recent interspecific gene flow. However, these admixed populations differentiated into specific genetic clusters when assuming eight genetic clusters. Contrary to expectations, the genetic distance between the eight genetic clusters does not clearly delineate species boundary, but instead shows a continuous gradient in genetic distance that suggests either ancient hybridization or retention of ancestral polymorphism. Notably, within-species genetic differentiation estimates of 0.028 and 0.047 for *Q. canariensis* and *Q. faginea* respectively, are intermediate between typical estimates of microsatellite genetic differentiation among European white oaks core range populations (Muir *et al*., 2004; Alberto *et al*., 2010; Neophytou *et al*., 2015) and isolated marginal populations (Buschbom *et al*., 2011; Moracho *et al*., 2016). Results herein therefore suggest that interspecific gene flow occurred before population divergence in isolation. Divergence by drift was found to be the driving factor shaping genetic structure of small isolated *Q. faginea* and *Q. canariensis* populations. Recent drift in isolation was exacerbated in declining *Q. faginea* populations and affected to a lesser extent the remaining *Q. faginea* and some *Q. canariensis* populations. This has led to the formation of independent genetic clusters in a context of constant effective population size, allowing them to retain some admixed characteristics from ancient hybridization events as shown by intermediate genetic characteristics of their respective genetic clusters. Indeed, populations that experienced stable environment for extended periods of time are expected to retain ancestral characteristics (Petit *et al*., 2005), and thus in our context, the footprint of ancient interspecific gene flow. As a result of the recent evolution by drift in isolation, each population constitutes an original and significant contribution to the total genetic diversity of the species and provides a good illustration that rear-edge populations effectively act as reservoirs of evolutionary history (Petit *et al*., 2005; Médail and Diadema, 2009; Lepais *et al*., 2013), as suggested by the description of subspecies in the region (Aissi *et al*., 2021). Given their small population size, these populations are vulnerable to rapid decline. This is supported by our simulations showing a demographic decline that has not yet led to a reduction in population genetic diversity. However, ABC-based estimates of historical declines in some *Q. faginea* populations was not always congruent with linkage-disequilibrium based estimates of small contemporary effective population size, thus indicating that all populations might not follow the same recent demographic trends. Nevertheless, most of the studied populations have effective population size confidence intervals below 500 individuals, a critical level for the long term maintenance of evolutionary potential (Jamieson and Allendorf, 2012; Hoban *et al*., 2020), and estimates of effective population sizes as low as 50 individuals in isolated *Q. faginea* populations suggest short-term risk of detrimental effect of inbreeding depression on population persistence (Jamieson and Allendorf, 2012). If these populations are probably more likely to be in immediate threat posed by demographic stochasticity rather than generic risks, long term retention of evolutionary potential is however paramount for adaptation to environmental changes (Allendorf and Luikart, 2007) and is clearly challenged by low effective population size of rear-edge *Q. faginea* populations. Additional studies at the species distribution range level, including core and marginal populations should provide a better understanding of the history and evolutionary significance of intraspecific genetic diversity within these Mediterranean oak species.

### The value of compound markers for population history reconstruction

Sequence information around microsatellite loci can uncover molecular variations caused by different mutation dynamics and mechanisms that generate complementary patterns across populations and species (Hey *et al*., 2004; Barthe *et al*., 2012). If the dataset combining all polymorphisms under microhaplotypes showed the most clear-cut results to depict the geographical distribution of the genetic clusters, we also observed different patterns of diversity distribution across populations according to the polymorphisms considered. For instance, microsatellite variation drives the build-up of genetic differentiation among *Q. faginea* populations, but not among *Q. canariensis* populations. Given that microsatellite loci evolve faster than DNA in their flanking sequences, increased differentiation at fast-evolving loci might suggest recent demographic events affecting *Q. faginea* populations. Geographic patterns of microsatellite differentiation are not linked to newly acquired mutations because accounting for phylogenetic distance between microsatellite alleles does not yield to higher population differentiation. This suggests that other factors, such as drift or interspecific gene flow, are involved in the increased differentiation at microsatellites for *Q. faginea*. However, highly polymorphic multi-allelic microsatellite should be more sensitive to the recent effect of genetic drift. Remarkably, estimates of genetic differentiation among *Q. faginea* populations were similar to estimates of genetic differentiation between *Q. faginea* and *Q. canariensis*, suggesting that interspecific gene flow blurs species boundaries with morphologically undetected hybridization and directional introgression introducing heterospecific alleles within *Q. faginea* populations. A combination of interspecific gene flow and recent demographic events could account for this pattern of molecular variation, highlighting the limits of estimates of genetic diversity and population genetic structure to resolve complex demographic history. Nevertheless, contrasted variability between differently evolving polymorphisms suggests complementary information about past population history. We showed that simulation-based inferences, that allow for realistic modelling of molecular evolution at linked loci in populations, leverage information complementarily for improved historical inferences.

We found contrasted most likely demographic scenarios when considering either microsatellites alone or in conjunction with variation in their flanking sequences. Recent interspecific gene flow was inferred when using microsatellite variation alone while divergence without interspecific gene flow was the most likely scenario when also accounting for flanking sequence variation. Homoplasy (identical allelic state derived from different ancestor alleles) at microsatellite loci explains this discrepancy: with the underestimation of interspecific divergence due to homoplasy (Estoup *et al*., 2002) interspecific gene flow needs to be invoked to explain the observed pattern of microsatellite variation between species. Indeed, homoplasy at microsatellite loci is expected to overestimate the estimation of hybridization rate and underestimates the estimation of the timing of interspecific gene flow events (Dickey *et al*., 2013; Henriques *et al*., 2016).

However, contrary to traditional implementation of ABC, Random Forest ABC is not sensitive to the number and relevance of the summary statistics (Marin *et al*., 2018) and can assess the weight of each summary statistics in the inference (Estoup *et al*., 2018; Raynal *et al*., 2019). Therefore, we took advantage of theses new properties to compare the value of microsatellite variations, flanking sequence substitutions, or the combination of both, to inform biological inferences. For model choice, all types of polymorphisms provide some information to differentiate the concurrent demographic models. For population demographic parameters, population-level diversity and pairwise difference between populations computed from haplotype and microsatellites inform about effective population size of sampled populations. However, the level of genetic diversity within the flanking sequence brings some additional information for ancient events such as effective size of ancestral populations. While population-level correlation between genetic diversity at flanking sequence and microsatellite was suggested to be sensitive to past demographic events (Payseur and Cutter, 2006), nine out of 32 of those summary statistics were among the 33 uninformative summary statistics out of the total of 518 summary statistics derived in this study. This indicates that not all summary statistics are informative with respect to parameter inference, irrespectively of the type of molecular variability they summarized. Deeper assessment of correlation between diversity statistics (Gaggiotti *et al*., 2018) at contrasted polymorphisms or the direct use of raw haplotypes without summary with deep learning approaches (Flagel *et al*., 2019) seem promising avenues to extract more information from multi-polymorphisms marker sequences.

Last but not least, we found that microsatellite mutation rate at HapSTR and SSR loci and substitution rate at their flanking sequence are overall informed by population level summary statistics, while locus-specific mutation parameters are specifically informed by the corresponding locus-level summary statistics. These results confirm the relevance of inference at the locus level in order to capture a realistic picture of mutational dynamic variability across loci (Payseur and Cutter, 2006; Chapuis *et al*., 2020). Given the renewed interest in the functional role of microsatellite variation (Xie *et al*., 2019; Press *et al*., 2019) and the availability of novel ways to sequence thousands of targeted microsatellite loci in empirical populations (Press *et al*., 2018), locus-specific inference contrasting flanking sequence and microsatellite mutation rates could help identifying functionally relevant microsatellite variation showing unusually low or high rate of evolution. Integrating different sources of polymorphism will thus further our understanding of the role of molecular variations in shaping species adaptation.

## Acknowledgments

We thank Rémy Petit and Cecile Bacles for advice and comments on a previous version of the manuscript. Sequence-based microsatellite genotyping was performed at the PGTB (doi:10.15454/1.5572396583599417E12) with the help of Erwan Guichoux and Emilie Chancerel. Computer time for the demographic simulations and ABC inferences was provided by the computing facilities MCIA (Mésocentre de Calcul Intensif Aquitain) of the Université de Bordeaux and of the Université de Pau et des Pays de l’Adour.

## Author Contributions

AA, EV, YB and OL obtained the funding for the research, contributed to the design of the study and the interpretation of the results. AA performed the sampling and collected field data under the supervision of YB. AA and OL performed DNA extraction. OL performed the laboratory work to produce the sequencing data, the bioinformatic and population genetic data analysis, drafted the first version of the manuscript and handled the manuscript submission and revisions. All authors contributed to the article revisions and approved the final version.

## Data archiving

The datasets of *Q. faginea* and *Q. canariensis* genotypes used in this study will be made publicly available in Dryad upon manuscript acceptance.

## References

Aissi A, Beghami Y, Heuertz M (2019). Le chêne faginé (*Quercus faginea*, Fagaceae) en Algérie : potentiel germinatif et variabilité morphologique des glands et des semis. Plant Ecol Evol 152: 437–449.

Aissi A, Beghami Y, Lepais O, Véla E (2021). Morphological and taxonomic analysis of *Quercus faginea* (Fagaceae) complex in Algeria. Botany 99: 99–113.

Alberto F, Niort J, Derory J, Lepais O, Vitalis R, Galop D, et al. (2010). Population differentiation of sessile oak at the altitudinal front of migration in the French Pyrenees. Mol Ecol 19: 2626–2639.

Allendorf FW, Luikart G (2007). Conservation and the genetics of populations. Blackwell Pub.

De Barba M, Miquel C, Lobréaux S, Quenette PY, Swenson JE, Taberlet P (2016). High-throughput microsatellite genotyping in ecology: Improved accuracy, efficiency, standardization and success with low-quantity and degraded DNA. Mol Ecol Resour 17: 492–507.

Barthe S, Gugerli F, Barkley NA, Maggia L, Cardi C, Scotti I (2012). Always look on both sides: phylogenetic information conveyed by simple sequence repeat allele sequences. PLoS One 7: e40699.

Beaumont M a, Zhang W, Balding DJ (2002). Approximate Bayesian computation in population genetics. Genetics 162: 2025–35.

Belkhir K, Borsa P, Chikhi L, Raufaste N, Bonhomme F (2004). GENETIX 4.05, logiciel sous Windows TM pour la génétique des populations. Laboratoire Génome, Populations, Interactions, CNRS UMR 5171, Université de Montpellier II, Montpellier (France).

Bradbury IR, Wringe BF, Watson B, Paterson I, Horne J, Beiko R, et al. (2018). Genotyping-by-sequencing of genome-wide microsatellite loci reveals fine-scale harvest composition in a coastal Atlantic salmon fishery. Evol Appl 11: 918–930.

Buschbom J, Yanbaev Y, Degen B (2011). Efficient long-distance gene flow into an isolated relict oak stand. J Hered 102: 464–472.

Chapuis M, Raynal L, Plantamp C, Meynard CN, Blondin L, Marin J, et al. (2020). A young age of subspecific divergence in the desert locust inferred by ABC Random Forest. Mol Ecol 29: 4542–4558.

Cornuet J-M, Ravigné V, Estoup A (2010). Inference on population history and model checking using DNA sequence and microsatellite data with the software DIYABC (v1.0). BMC Bioinformatics 11: 401.

Curto M, Winter S, Seiter A, Schmid L, Scheicher K, Barthel LMF, et al. (2019). Application of a SSR-GBS marker system on investigation of European Hedgehog species and their hybrid zone dynamics. Ecol Evol 9: 2814–2832.

Darby BJ, Erickson SF, Hervey SD, Ellis-Felege SN (2016). Digital fragment analysis of short tandem repeats by high-throughput amplicon sequencing. Ecol Evol 6: 4502–4512.

Dickey AM, Hall PM, Shatters RG, Mckenzie CL (2013). Evolution and homoplasy at the Bem6 microsatellite locus in three sweetpotato whitefly (*Bemisia tabaci*) cryptic species. BMC Res Notes 6: 249.

Do C, Waples RS, Peel D, Macbeth GM, Tillett BJ, Ovenden JR (2013). NeEstimator v2: re-implementation of software for the estimation of contemporary effective population size (Ne) from genetic data. Mol Ecol Resour 14: 209–214.

Durand J, Bodenes C, Chancerel E, Frigerio JM, Vendramin G, Sebastiani F, et al. (2010). A fast and cost-effective approach to develop and map EST-SSR markers: oak as a case study. BMC Genomics 11: 570.

Earl DA, vonHoldt BM (2012). STRUCTURE HARVESTER: a website and program for visualizing STRUCTURE output and implementing the Evanno method. Conserv Genet Resour 4: 359–361.

Estoup A, Jarne P, Cornuet J-M (2002). Homoplasy and mutation model at microsatellite loci and their consequences for population genetics analysis. Mol Ecol 11: 1591–1604.

Estoup A, Raynal L, Verdu P, Marin J-M (2018). Model choice using Approximate Bayesian Computation and Random Forests: analyses based on model grouping to make inferences about the genetic history of Pygmy human populations. J la Société Fr Stat 159: 167–190.

Excoffier L, Dupanloup I, Huerta-Sánchez E, Sousa VC, Foll M (2013). Robust demographic inference from genomic and SNP data. PLoS Genet 9: e1003905.

Excoffier L, Lischer HEL (2010). Arlequin suite ver 3.5: a new series of programs to perform population genetics analyses under Linux and Windows. Mol Ecol Resour 10: 564–567.

Falush D, Stephens M, Pritchard JK (2003). Inference of population structure using multilocus genotype data: linked loci and correlated allele frequencies. Genetics 164: 1567–87.

Feliner GN (2014). Patterns and processes in plant phylogeography in the Mediterranean Basin. A review. Perspect Plant Ecol, Evol Syst 16: 265–278.

Flagel L, Brandvain Y, Schrider DR (2019). The unreasonable effectiveness of convolutional neural networks in population genetic inference. Mol Biol Evol 36: 220–238.

Foll M, Gaggiotti O (2008). A genome-scan method to identify selected loci appropriate for both dominant and codominant markers: A Bayesian perspective. Genetics 180: 977–993.

Gaggiotti OE, Chao A, Peres-Neto P, Chiu CH, Edwards C, Fortin MJ, et al. (2018). Diversity from genes to ecosystems: A unifying framework to study variation across biological metrics and scales. Evol Appl 11: 1176–1193.

García Murillo P., Harvey-Brown Y (2017). Quercus canariensis. In: The IUCN Red List of Threatened Species,, p e.T78809256A80570536.

Gómez A, Lunt DH (2007). Refugia within refugia: Patterns of phylogeographic concordance in the Iberian peninsula. In: Weiss S, Ferrand N (eds) Phylogeography of Southern European Refugia, Springer Netherlands: Dordrecht, pp 155–188.

Gorener V, Harvey-Brown Y, Barstow M (2017). Quercus canariensis. IUCN red List Threat species e.T7880925.

Goudet J (1995). FSTAT (Version 1.2): A computer program to calculate F-statistics. J Hered 86: 485–486.

Hampe A, Petit RJ (2005). Conserving biodiversity under climate change: the rear edge matters. Ecol Lett 8: 461–7.

Hardy OJ, Vekemans X (2002). SPAGeDi: a versatile computer program to analyse spatial genetic structure at the individual or population levels. Mol Ecol Notes 2: 618–620.

Harvey-Brown Y, García Murillo PG, Buira A (2017). Quercus faginea. IUCN Red List Threat Species: e.T78916251A80570540.

Henriques R, von der Heyden S, Matthee CA (2016). When homoplasy mimics hybridization: a case study of Cape hakes (*Merluccius capensis* and *M. paradoxus*). PeerJ 4: e1827.

Hewitt GM (1999). Post-glacial re-colonization of European biota. Biol J Linn Soc 68: 87–112.

Hey J, Won YJ, Sivasundar A, Nielsen R, Markert JA (2004). Using nuclear haplotypes with microsatellites to study gene flow between recently separated Cichlid species. Mol Ecol 13: 909–919.

Hoban S, Bruford M, D’Urban Jackson J, Lopes-Fernandes M, Heuertz M, Hohenlohe PA, et al. (2020). Genetic diversity targets and indicators in the CBD post-2020 Global Biodiversity Framework must be improved. Biol Conserv 248: 108654.

Hoogenboom J, de Knijff P, Laros JFJ, de Leeuw RH, van der Gaag KJ, Sijen T (2016). FDSTools: A software package for analysis of massively parallel sequencing data with the ability to recognise and correct STR stutter and other PCR or sequencing noise. Forensic Sci Int Genet 27: 27–40.

Jamieson IG, Allendorf FW (2012). How does the 50/500 rule apply to MVPs? Trends Ecol Evol 27: 578–584.

Jerome D, Vasquez F (2018). Quercus faginea. IUCN Red List Threat Species e.T7891625.

Kalinowski ST (2005). HP-RARE 1.0: A computer program for performing rarefaction on measures of allelic richness. Mol Ecol Notes 5: 187–189.

Kampfer S, Lexer C, Glössl J, Steinkellner H (1998). Characterization of (GA)n microsatellite loci from *Quercus robur*. Hereditas 129: 183–186.

Kivelä M, Arnaud-Haond S, Saramäki J (2015). EDENetworks: A user-friendly software to build and analyse networks in biogeography, ecology and population genetics. Mol Ecol Resour 15: 117–122.

Kopelman NM, Mayzel J, Jakobsson M, Rosenberg NA, Mayrose I (2015). Clumpak: A program for identifying clustering modes and packaging population structure inferences across K. Mol Ecol Resour 15:1179–1191.

Layton KKS, Dempson B, Snelgrove PVR, Duffy SJ, Messmer AM, Paterson IG, et al. (2020). Resolving fine-scale population structure and fishery exploitation using sequenced microsatellites in a northern fish. Evol Appl: eva.12922.

Lepais O, Chancerel E, Boury C, Salin F, Manicki A, Taillebois L, et al. (2020). Fast sequence-based microsatellite genotyping development workflow. PeerJ 8: e9085.

Lepais O, Leger V, Gerber S (2006). Short note: high throughput microsatellite genotyping in oak species. Silvae Genet 55: 238.

Lepais O, Muller SD, Ben Saad-Limam S, Benslama M, Rhazi L, Belouahem-Abed D, et al. (2013). High genetic diversity and distinctiveness of rear-edge climate relicts maintained by ancient tetraploidisation for Alnus glutinosa. PLoS One 8: e75029.

Leroy T, Roux C, Villate L, Bodénès C, Romiguier J, Paiva JAP, et al. (2017). Extensive recent secondary contacts between four European white oak species. New Phytol 214: 865–878.

Lye GC, Lepais O, Goulson D (2011). Reconstructing demographic events from population genetic data: the introduction of bumblebees to New Zealand. Mol Ecol 20: 2888–900.

Magri D, Fineschi S, Bellarosa R, Buonamici A, Sebastiani F, Schirone B, et al. (2007). The distribution of *Quercus suber* chloroplast haplotypes matches the palaeogeographical history of the western Mediterranean. Mol Ecol 16: 5259–66.

Marin J, Pudlo P, Estoup A, Robert C (2018). Likelihood-free model choice. In: Sisson S a, Fan Y, Beaumont M (eds) Handbook of Approximate Bayesian Computation, CRC Press, p 153.

Médail F, Diadema K (2009). Glacial refugia influence plant diversity patterns in the Mediterranean Basin. J Biogeogr 36: 1333–1345.

Moracho E, Moreno G, Jordano P, Hampe A (2016). Unusually limited pollen dispersal and connectivity of Pedunculate oak (*Quercus robur*) refugial populations at the species’ southern range margin. Mol Ecol 25: 3319–3331.

Mountain JL, Knight A, Jobin M, Gignoux C, Miller A, Lin AA, et al. (2002). SNPSTRs: Empirically derived, rapidly typed, autosomal haplotypes for inference of population history and mutational processes. Genome Res 12: 1766–1772.

Muir G, Lowe AJ, Fleming CC, Vogl C (2004). High nuclear genetic diversity, high levels of outcrossing and low differentiation among remnant populations of *Quercus petraea* at the margin of its range in Ireland. Ann Bot 93: 691–697.

Neophytou C, Gärtner SM, Vargas-Gaete R, Michiels H-G (2015). Genetic variation of Central European oaks: shaped by evolutionary factors and human intervention? Tree Genet Genomes 11: 1–15.

Payseur BA, Cutter AD (2006). Integrating patterns of polymorphism at SNPs and STRs. Trends Genet 22: 424–429.

Petit RJ, Brewer S, Bordács S, Burg K, Cheddadi R, Coart E, et al. (2002). Identification of refugia and post-glacial colonisation routes of European white oaks based on chloroplast DNA and fossil pollen evidence. For Ecol Manage 156: 49–74.

Petit RJ, Hampe A, Cheddadi R (2005). Climate changes and tree phylogeography in the Mediterranean. Taxon 54: 877–885.

Press MO, Hall AN, Morton EA, Queitsch C (2019). Substitutions are boring: Some arguments about parallel mutations and high mutation rates. Trends Genet 35: 253–264.

Press MO, Mccoy RC, Hall AN, Akey JM, Queitsch C (2018). Massive variation of short tandem repeats with functional consequences across strains of *Arabidopsis thaliana*. Genome Res 28: 1169–1178.

Pritchard JK, Stephens M, Donnelly P (2000). Inference of population structure using multilocus genotype data. Genetics 155: 945–59.

Pudlo P, Marin J-M, Estoup A, Cornuet J-M, Gautier M, Robert CP (2016). Reliable ABC model choice via random forests. Bioinformatics 32: 859–866.

Ramakrishnan U, Mountain JL (2004). Precision and accuracy of divergence time estimates from STR and SNPSTR variation. Mol Biol Evol 21: 1960–1971.

Raynal L, Marin J-M, Pudlo P, Ribatet M, Robert CP, Estoup A (2019). ABC random forests for Bayesian parameter inference. Bioinformatics 35: 1720–1728.

Rodríguez-Sánchez F, Hampe A, Jordano P, Arroyo J (2010). Past tree range dynamics in the Iberian Peninsula inferred through phylogeography and palaeodistribution modelling: A review. Rev Palaeobot Palynol 162: 507–521.

Šarhanová P, Pfanzelt S, Brandt R, Himmelbach A, Blattner FR (2018). SSR-seq: Genotyping of microsatellites using next-generation sequencing reveals higher level of polymorphism as compared to traditional fragment size scoring. Ecol Evol 8: 10817–10833.

Scotti-Saintagne C, Mariette S, Porth I, Goicoechea PG, Barreneche T, Bodénès C, et al. (2004). Genome scanning for interspecific differentiation between two closely related oak species *[Quercus robur* L. and *Q. petraea* (Matt.) Liebl.]. Genetics 168: 1615–26.

Vartia S, Villanueva-Cañas JL, Finarelli J, Farrell ED, Collins PC, Hughes GM, et al. (2016). A novel method of microsatellite genotyping-by-sequencing using individual combinatorial barcoding. R Soc Open Sci 3: 150565.

Viruel J, Haguenauer A, Juin M, Mirleau F, Bouteiller D, Boudagher-Kharrat M, et al. (2018). Advances in genotyping microsatellite markers through sequencing and consequences of scoring methods for *Ceratonia siliqua* (Leguminosae). Appl Plant Sci 6: e01201.

Wang J (2016). Individual identification from genetic marker data: developments and accuracy comparisons of methods. Mol Ecol Resour 16: 163–175.

Waples RS, Do C (2010). Linkage disequilibrium estimates of contemporary Ne using highly variable genetic markers: A largely untapped resource for applied conservation and evolution. Evol Appl 3: 244–262.

Xie KT, Wang G, Thompson AC, Wucherpfennig JI, Reimchen TE, MacColl ADC, et al. (2019). DNA fragility in the parallel evolution of pelvic reduction in stickleback fish. Science (80-) 363: 81–84.

