## Supporting Material File 1 for "Joint analysis of microsatellites and flanking sequences enlightens complex demographic history of interspecific gene flow and vicariance in rear-edge oak populations"

**Supporting Material and Methods**

**Sampling**

**Table S1**: Description of the sampled *Q. faginea* and *Q. canariensis* Algerian population.

| Site | Taxonomic identity | Location | Latitude | Longitude | Number of sampled individuals | Proportion of standing individual sampled |
| --- | --- | --- | --- | --- | --- | --- |
| S01 | *Q. canariensis* | Hafir Tlemcen | 34.8326 | -1.3748 | 30 | 5% |
| S02 | *Q. faginea* | Terni Tlemcen | 34.7772 | -1.3440 | 30 | 40% |
| S03 | *Q. faginea* | Baloul Saïda | 34.9844 | 0.4041 | 30 | 60% |
| S08 | *Q. faginea* | Safalou Tiaret | 35.4053 | 1.3486 | 30 | 10% |
| S07 | *Q. canariensis* | Thniet El Hed Tessemssilt | 35.8655 | 1.9776 | 30 | 5% |
| S06 | *Q. canariensis* | Errich Bouira | 36.4057 | 3.8664 | 30 | 5% |
| S10 | *Q. canariensis* | Akfadou Tizi-Ouzou | 36.7218 | 4.4546 | 29 | <1% |
| S09 | *Q. canariensis* | Hamza Jijel | 36.7483 | 5.6065 | 28 | <1% |
| S11 | *Q. faginea* | Chélia Aurés | 35.3679 | 6.6320 | 28 | 5% |
| S04 | *Q. canariensis* | Mechrouha Souk Ahras | 36.3856 | 7.8606 | 30 | <1% |
| S05 | *Q. canariensis* | Ghora Bougous El-Taref | 36.6739 | 8.4702 | 29 | <1% |

**Microsatellite genotyping by sequencing**

DNA from each individual was extracted using Invisorb DNA Plant HTS 96 Kit (Invitek) from 35 mg of dried leaf following manufacturer instructions and checked by 1.5% agarose gel electrophoresis. Sequence-based microsatellite genotyping (SSRseq) was used to analyse polymorphism at 36 loci as described elsewhere (Lepais et al., 2020). In short, 60 genomic and EST-derived microsatellites (Kampfer et al., 1998; Durand et al., 2010) were co-amplified in a single multiplexed PCR using redesigned primers to target fragments between 120 and 200 bp, including Nextera Illumina tags at 5’ ends of primers. Following two round of locus specific amplification, each of the 384 samples (330 individuals in addition to 48 repeated individuals, two negative controls (water instead of DNA) and two *Quercus robur* and two *Quercus petraea* as additional controls) was indexed using one of the 384 barcode combination from the Nextera XT index set. Amplicons from each of the four 96-well plates were then pooled, purified with 1.8X Agencourt AMPure XP beads (Beckman Coulter), quality checked using Agilent Tapestation D1000 kit and Qubit fluorometric system (Thermo Fisher Scientific) and quantified using Kapa library quantification kit in a Roche LightCycler 480 quantitative PCR. The resulting four pools were then pooled in equimolar concentration and sequenced using 1/3 of an Illumina MiSeq v2 flowcell 2x250 bp paired-end sequencing kit. After sequence demultiplexing, a bioinformatics pipeline (Lepais et al., 2020) integrating the FDSTools analysis toolkit (Hoogenboom et al., 2016) was used to convert raw sequence into genotypic data (microhaplotypes) integrating all polymorphisms identified within the microsatellite itself (number of repeats, SNP and Indel) and in the flanking sequences (SNP and Indel). The bioinformatics pipeline also compares genotypes from repeated individuals to estimate allelic error rate and compute the overall missing data rate for each locus. This qualitative information allows optimising the bioinformatics analysis strategy, in particular identify loci that cannot be reliably genotyped or for which the genotyping accuracy is only acceptable when analysing the polymorphism within the repeat motif itself, i.e. not accounting for the polymorphism in the flanking sequence (Lepais et al., 2020). Finally, for each locus, every allele differing from the other by any polymorphism was coded under an arbitrary three-digit scheme with a unique number assigned to unique to each microhaplotype within each locus.

**Population demographic history**

**Prior distribution** of demographic parameters has been setup wide to reflect knowledge uncertainty in the history of these populations. Effective population sizes (*N*) were parameterized as log uniform distribution between 10 and 1,000,000 haploid genomes. Variation of effective population size (expansion or contraction) was account for using a growth rate parameter (*g*) that linked current and ancestral (at time of divergence, *Na*) effective population sizes as well as divergence time of studied populations (Figure 3, *t*). Divergence time between core range of *Q. faginea* and *Q. canariensis* (*ta*) as uniform prior distribution between 401 and 10,000 generations (corresponding to 20,050 to 500,000 years assuming a generation time of 50 years (Gregorius, Degen & König, 2007; Leroy et al., 2020)) and divergence time between core range and studied population (*t*) between 10 and 400 generations (i.e. 500 to 20,000 years). As contemporary and ancestral effective population size had the same prior definition, a wide range of exponential population dynamics were considered (stable, growth or decline). Note that the core range ghost populations (*N8* and *N9*) and their ancestral population (*N10*) were considered fixed with value between 10 and 1,000,000 following a log-uniform distribution. Bidirectional interspecific gene flow between species core range ghost populations (*M89* and *M98*) were setup as uniform between 0.0001 and 0.01 haploid gametes per generation. The rate of directional gene flow between heterospecific core range and studied populations (*M*) followed a uniform distribution between 0.0001 and 0.01 haploid gametes per generation. The proportion of migrant during instantaneous event of interspecific gene flow (*R*) as a uniform distribution between 0.05 and 0.95. Interspecific gene influx into the studied population happened after an initial phase of isolation (*tx*, uniform between 5 and 200 generations) at a time (episodic event) or since a time (continuous gene flow) between 5 and 200 generations (Figure 3). Prior range have been setup in a way that model should differ enough to result into different level of genetic diversity and structure and being identifiable from each other. For instance, when gene flow is allowed in a model it should be significant, otherwise model with or without gene flow will be undistinguishable. Prior ranges have been checked for their capacity to produce realistic genotypic data by comparing the distribution of each summary statistics (see below) computed from the simulated dataset with those of the observed dataset (Cornuet, Ravigné & Estoup, 2010; Leroy et al., 2017a; Supporting Material File 2). Early simulations attempts using sub-optimal prior range definitions resulted in most of the observed summary statistics to fall outside the 99% quantile of the simulated summary statistic distributions, a clear indication that simulations were not able to produce realistic genotypic dataset.

**Supporting Results**

**Genotyping**

Form the 60 loci included in the multiplexed PCR, 53 produced at least 20 sequences in more than 50% of the individuals and were subsequently analysed. Of these, 39 loci with less than 15% of missing genotypes (1.5% of missing genotypes across the whole dataset) and 6% of allelic error (0.41% of overall allelic error) were kept for further inspection (Table S1). Following testing for Hardy-Weinberg equilibrium, 3 additional loci that show significant deficit in heterozygotes were removed. Only QrZAG112 showed significant but moderate signal of balancing selection when analysed to the interspecific level (qval=0.056) but not at the intraspecific level (qval > 0.10; Supporting Material File 2). We therefor assumed that the 36 remaining loci behaves as expected under neutrally.

Following the removal of 6 individuals that failed to amplify or showed more than 4 missing genotypes and of additional 6 individuals that happened to represent ramets of the same genet, showing anecdotic clonality events, 318 individuals remained for subsequent analyses.

Table S2: Characteristics of the studied genetic markers.

| Locus | Origin | Type of locus | Missing data | Allelic error rate^a^ | Mean length^b^ | Microsatellite repeat | | | | Flanking sequence | | N allele size^d^ | N haplotype ^e^ | Size homoplasy |
| --- | --- | --- | --- | --- | --- | --- | --- | --- | --- | --- | --- | --- | --- | --- |
|  |  |  |  |  |  | SSR^c^ | Motif size | SNP | Indel | SNP^c^ | Indel |  |  |  |
| QpZAG15 | genomics | HapSTR | 0.00% | <1.0% | 87 | 1 | 2 | 0 | 0 | 4 | 0 | 10 | 19 | 47% |
| QrZAG74 | genomics | HapSTR | 0.00% | <1.0% | 68 | 1 | 2 | 1 | 0 | 10 | 1 | 3 | 18 | 83% |
| QrZAG96 | genomics | HapSTR | 0.00% | 1.0% | 75 | 1 | 2 | 0 | 0 | 5 | 2 | 19 | 22 | 14% |
| PIE022 | EST-based | HapSTR | 1.85% | <1.0% | 85 | 1 | 2 | 2 | 0 | 7 | 0 | 12 | 26 | 54% |
| PIE039 | EST-based | HapSTR | 0.00% | <1.0% | 77 | 1 | 3 | 1 | 1 | 8 | 2 | 9 | 17 | 47% |
| PIE102 | EST-based | HapSTR | 0.00% | 1.1% | 124 | 1 | 2 | 3 | 0 | 6 | 1 | 14 | 40 | 65% |
| PIE172 | EST-based | HapSTR | 0.00% | <1.0% | 118 | 1 | 6 | 3 | 0 | 11 | 0 | 6 | 14 | 57% |
| PIE187 | EST-based | HapSTR | 0.00% | <1.0% | 81 | 3 | 3; 3; 3 | 4 | 0 | 7 | 1 | 8 | 41 | 80% |
| PIE192 | EST-based | HapSTR | 0.31% | <1.0% | 103 | 1 | 3 | 0 | 0 | 5 | 0 | 6 | 11 | 45% |
| PIE196 | EST-based | HapSTR | 0.31% | <1.0% | 137 | 2 | 3; 3 | 6 | 0 | 12 | 0 | 5 | 20 | 75% |
| PIE198 | EST-based | HapSTR | 0.31% | <1.0% | 93 | 2 | 3; 3 | 7 | 0 | 9 | 0 | 5 | 18 | 72% |
| PIE200 | EST-based | HapSTR | 0.31% | <1.0% | 134 | 3 | 3; 3; 3 | 13 | 0 | 14 | 1 | 4 | 52 | 92% |
| PIE215 | EST-based | HapSTR | 0.00% | <1.0% | 67 | 1 | 3 | 1 | 0 | 3 | 0 | 8 | 11 | 27% |
| PIE227 | EST-based | HapSTR | 0.00% | <1.0% | 72 | 1 | 3 | 0 | 0 | 4 | 0 | 9 | 13 | 31% |
| PIE233 | EST-based | HapSTR | 0.00% | <1.0% | 100 | 1 | 3 | 3 | 0 | 8 | 0 | 10 | 21 | 52% |
| PIE246 | EST-based | HapSTR | 0.00% | <1.0% | 78 | 1 | 2 | 1 | 0 | 10 | 3 | 16 | 25 | 36% |
| PIE247 | EST-based | HapSTR | 0.31% | <1.0% | 69 | 1 | 2 | 2 | 0 | 8 | 4 | 19 | 49 | 61% |
| PIE248 | EST-based | HapSTR | 0.00% | <1.0% | 94 | 1 | 2 | 4 | 0 | 4 | 1 | 12 | 33 | 64% |
| PIE270 | EST-based | HapSTR | 0.00% | <1.0% | 36 | 1 | 2 | 3 | 0 | 9 | 2 | 12 | 23 | 48% |
| PIE273 | EST-based | HapSTR | 0.00% | <1.0% | 75 | 2 | 2; 2 | 3 | 0 | 6 | 0 | 13 | 35 | 63% |
| QrZAG112 | genomics | SSR | 0.00% | <1.0% | 21 | 1 | 2 | 2 | 0 | - | - | 9 | 13 | 31% |
| QrZAG11 | genomics | SSR | 0.00% | <1.0% | 32 | 1 | 2 | 2 | 0 | - | - | 9 | 10 | 10% |
| QpZAG1/5 | genomics | SSR | 1.85% | <1.0% | 31 | 2 | 2; 2 | 7 | 0 | - | - | 21 | 57 | 63% |
| QpZAG110 | genomics | SSR | 2.47% | 5.3% | 33 | 1 | 2 | 3 | 1 | - | - | 23 | 29 | 21% |
| PIE171 | EST-based | SSR | 0.93% | 1.0% | 32 | 2 | 2; 2 | 7 | 1 | - | - | 22 | 49 | 55% |
| PIE236 | EST-based | SSR | 0.00% | <1.0% | 37 | 2 | 3; 3 | 2 | 1 | - | - | 9 | 21 | 57% |
| PIE253 | EST-based | SSR | 1.54% | <1.0% | 22 | 1 | 2 | 5 | 2 | - | - | 17 | 20 | 15% |
| PIE271 | EST-based | SSR | 0.00% | <1.0% | 23 | 1 | 2 | 3 | 0 | - | - | 15 | 18 | 17% |
| PIE055 | EST-based | SSR | 0.00% | <1.0% | 22 | 1 | 2 | 3 | 1 | - | - | 8 | 13 | 38% |
| PIE100 | EST-based | SSR | 0.31% | <1.0% | 20 | 1 | 2 | 1 | 1 | - | - | 13 | 15 | 13% |
| PIE222 | EST-based | SSR | 1.85% | <1.0% | 18 | 1 | 3 | 4 | 0 | - | - | 5 | 7 | 29% |
| PIE095 | EST-based | SSR | 0.00% | 1.1% | 26 | 1 | 2 | 5 | 1 | - | - | 21 | 27 | 22% |
| PIE255 | EST-based | SSR | 4.63% | 1.1% | 33 | 1 | 2 | 7 | 1 | - | - | 27 | 61 | 56% |
| PIE267 | EST-based | SSR | 0.00% | <1.0% | 44 | 3 | 2; 2; 2 | 2 | 0 | - | - | 13 | 20 | 35% |
| PIE072 | EST-based | SSR | 0.00% | 1.0% | 24 | 1 | 2 | 3 | 0 | - | - | 13 | 16 | 19% |
| PIE141 | EST-based | SSR | 0.31% | 2.1% | 27 | 1 | 2 | 2 | 0 | - | - | 13 | 15 | 13% |
| Total |  |  | 0.48% | 0.4% |  |  |  |  |  |  |  | 438 | 899 | 51% |

^a^ computed from a blind-repeat genotyping of 48 diploids individuals (no allele mismatch observed means less than 1 error for 96 observations) ; ^b^ length of the DNA sequence analysed ; ^c^ polymorphisms considered in coalescent-based simulations; ^d^ number of alleles differing in length ; ^e^ number of allele integrating all observed polymorphism.

**Contemporary effective population size**

**Figure S1**: Contemporary effective population size estimated from the linkage disequilibrium methods for *Q. faginea* (S02, S03, S08, S11) and *Q. canariensis* (S01, S04, S05, S06, S07, S09, S10) Algerian populations. Point represents the mean and the bar shows 95% confidence interval. Population are ordered from west to east along the x-axis, colours refer to Structure results, horizontal dashed lines represent the average value across populations and vertical dashed lines uncertain effective population size estimates.

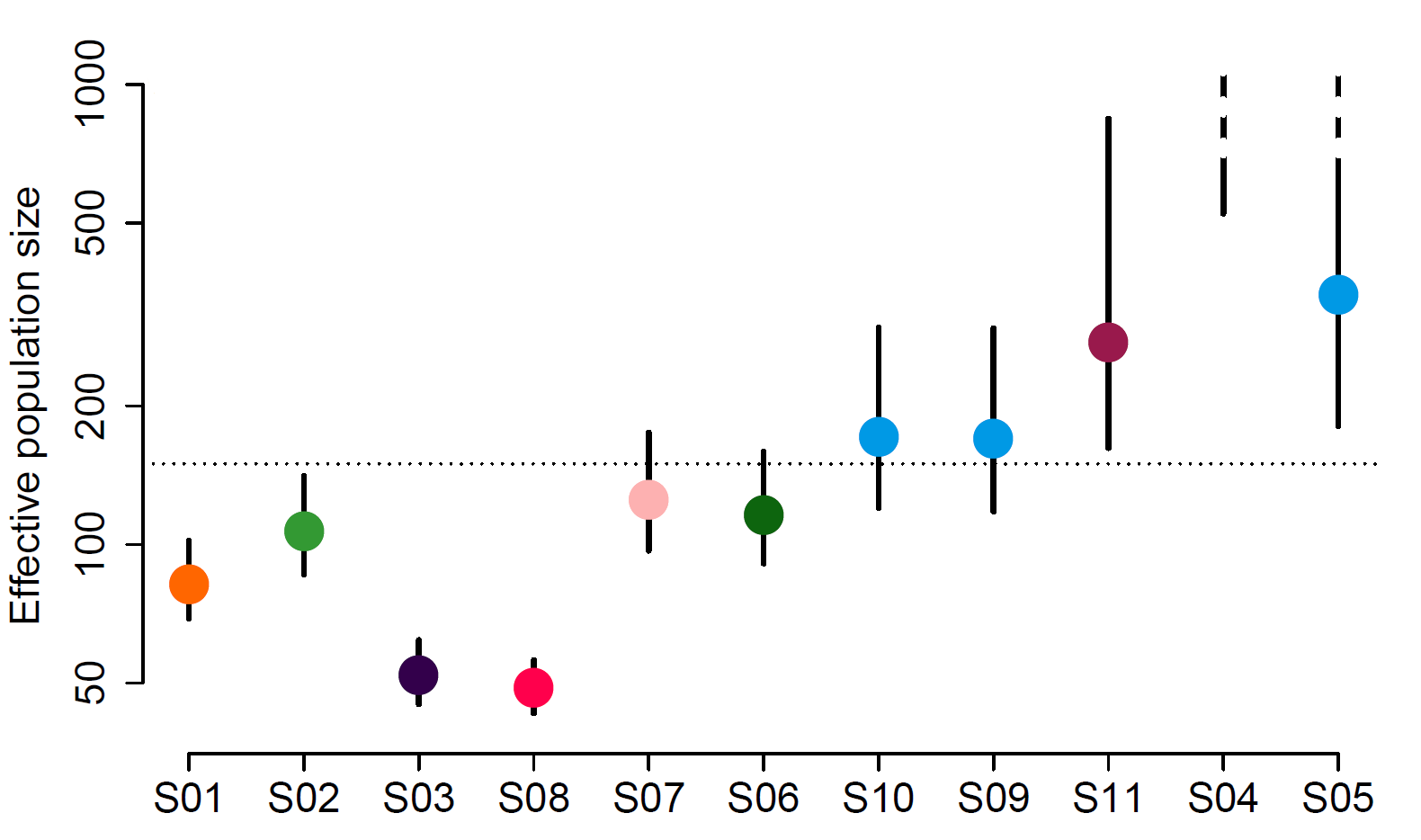

**Figure S2: Structure results across K value for the whole haplotype dataset**

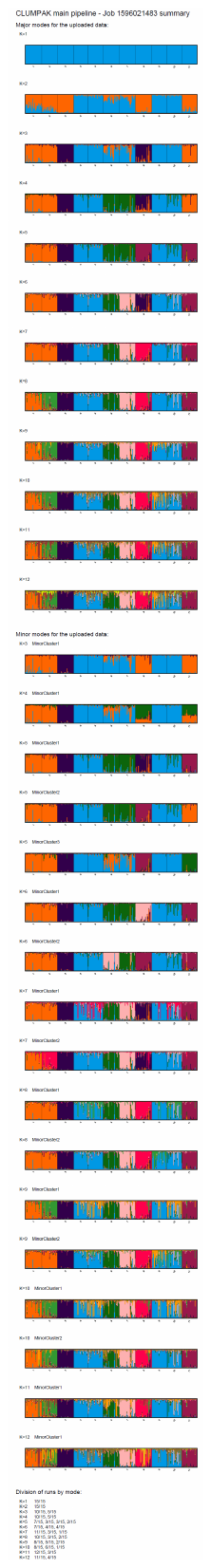

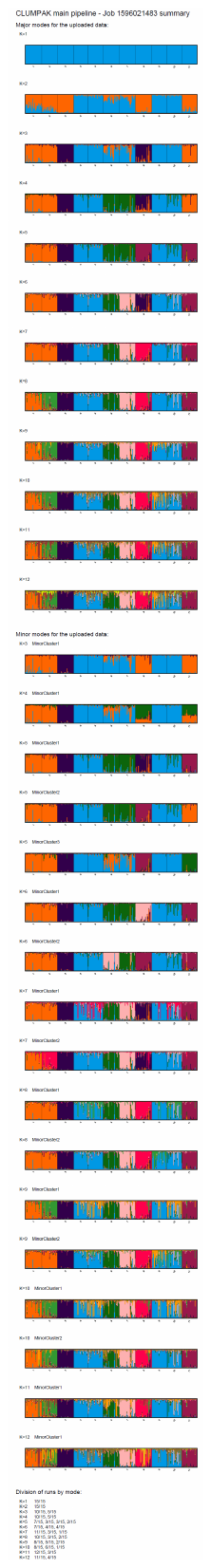

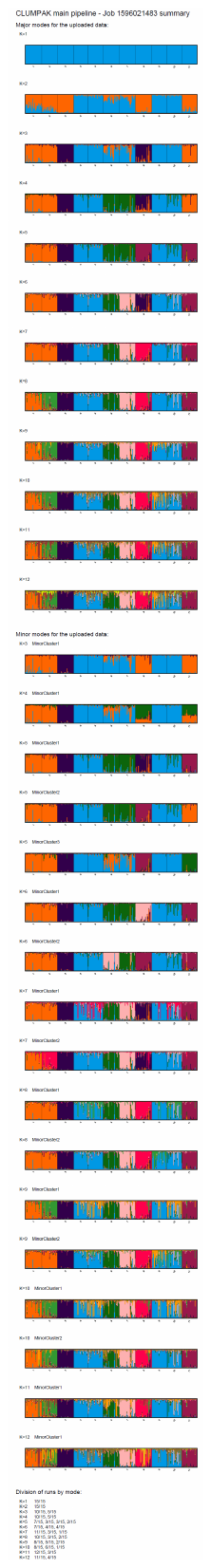

**Figure S3: Comparison of Structure results at k=2 and k=8 for different way to consider polymorphism.**

Haplotype (same results as manuscript Figure 2):

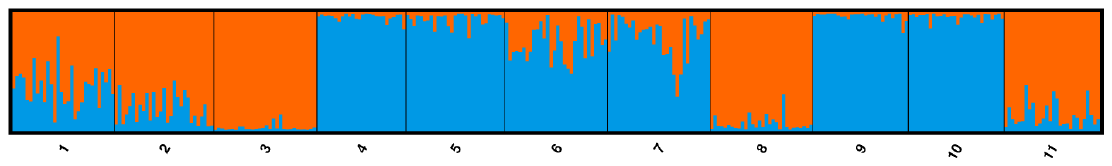

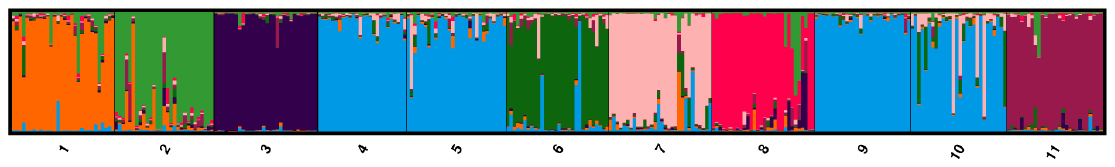

Number of microsatellite repeats:

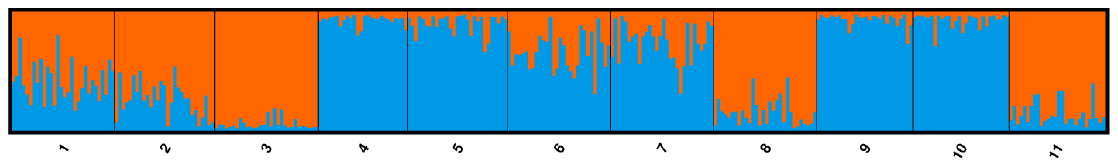

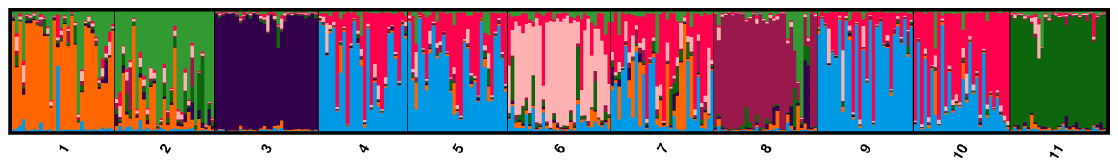

Allele size:

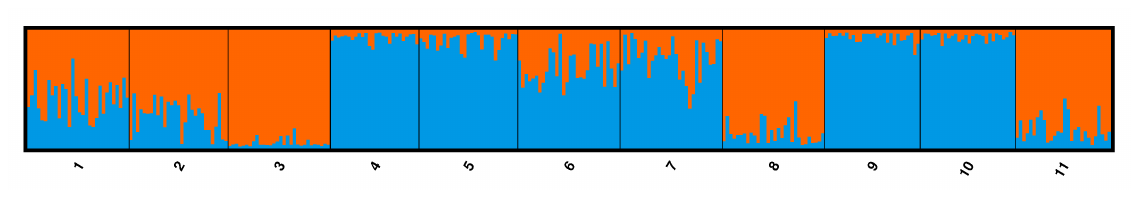

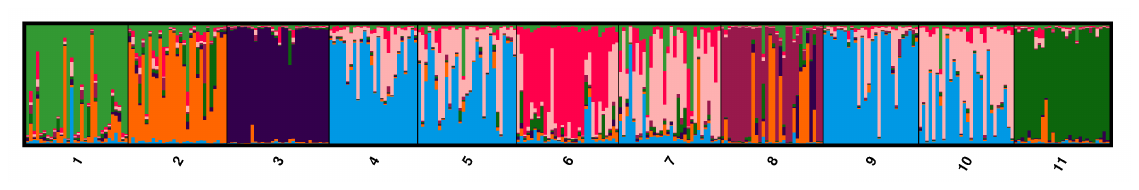

Flanking sequence haplotype:

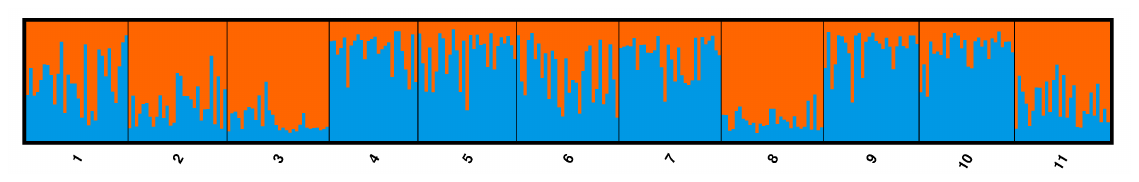

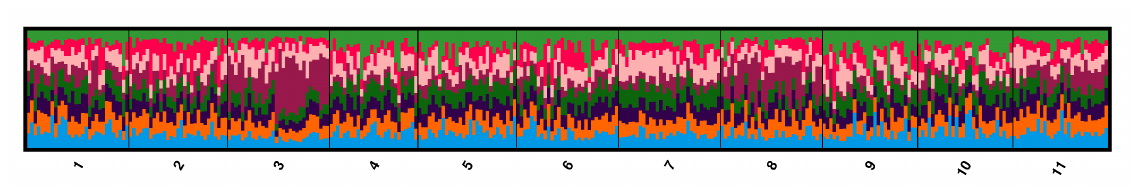

**Demographic inferences – summary statistics contribution**

The timing of events and level of interspecific gene flow were not estimable has showed by the posterior distribution that covers the whole of prior distribution (Table 4). Moreover, the number of informative summary statistics for these parameters were moderate (respectively, 21 and 18 on average for each parameter) and represented very low percentage of the total variable importance (13% and 15% respectively) suggesting a low signal-to-noise ratio for these parameters (Figure 4d et 4e).

For HapSTR loci, informative summary statistics to estimate microsatellite mutation rates originated at 61% from microsatellite variation, 37% from haplotype information and only 2% from the flanking sequence. As expected for SSR loci, most of the informative summary statistics originated from the microsatellite variation (91%) with information from haplotype and flanking sequence originating from other HapSTR loci only representing 6% and 2% respectively (Figure 4i). Finally, substitution rate at HapSTR loci were mostly derived from summary statistics composted from the flanking sequence substitutions (67% of the total variable importance of informative loci) followed by haplotype information (24%) and last microsatellite variation (9%).

For contemporary and ancestral effective population size of sampled populations, summary statistics computed from the microsatellite variation and from the haplotype contributed equally to the estimation (54% and 46%, respectively), while the flanking sequence substitutions did not contribute (0.2%). For effective population size of unsampled populations (i.e. *Q. canariensis* and *Q. faginea* core range, and their ancestor), summary statistics computed from the haplotype contributed for 59%, flanking sequence substitution for 31% and microsatellite variation a low 10% (figure 4c).

**Bibliography**

Cornuet J-M, Ravigné V, Estoup A. 2010. Inference on population history and model checking using DNA sequence and microsatellite data with the software DIYABC (v1.0). *BMC bioinformatics* 11:401. DOI: 10.1186/1471-2105-11-401.

Durand J, Bodenes C, Chancerel E, Frigerio JM, Vendramin G, Sebastiani F, Buonamici A, Gailing O, Koelewijn HP, Villani F, Mattioni C, Cherubini M, Goicoechea P, Herran A, Ikaran Z, Cabane C, Ueno S, Alberto F, Dumoulin PY, Guichoux E, Daruvar A de, Kremer A, Plomion C. 2010. A fast and cost-effective approach to develop and map EST-SSR markers: oak as a case study. *BMC Genomics* 11. DOI: 10.1186/1471-2164-11-570.

Gregorius H-R, Degen B, König A. 2007. Problems in the analysis of genetic differentiation among populations – a case study in Quercus robur. *Silvae Genetica* 56:190–199. DOI: 10.1515/sg-2007-0029.

Hoogenboom J, de Knijff P, Laros JFJ, de Leeuw RH, van der Gaag KJ, Sijen T. 2016. FDSTools: A software package for analysis of massively parallel sequencing data with the ability to recognise and correct STR stutter and other PCR or sequencing noise. *Forensic Science International: Genetics* 27:27–40. DOI: 10.1016/j.fsigen.2016.11.007.

Kampfer S, Lexer C, Glössl J, Steinkellner H. 1998. Characterization of (GA)n microsatellite loci from Quercus robur. *Hereditas* 129:183–186. DOI: 10.1111/j.1601-5223.1998.00183.x.

Lepais O, Chancerel E, Boury C, Salin F, Manicki A, Taillebois L, Dutech C, Aissi A, Bacles CFE, Daverat F, Launey S, Guichoux E. 2020. Fast sequence-based microsatellite genotyping development workflow. *PeerJ* 8:e9085. DOI: 10.7717/peerj.9085.

Leroy T, Rougemont Q, Dupouey J, Bodénès C, Lalanne C, Belser C, Labadie K, Le Provost G, Aury J, Kremer A, Plomion C. 2020. Massive postglacial gene flow between European white oaks uncovered genes underlying species barriers. *New Phytologist* 226:1183–1197. DOI: 10.1111/nph.16039.

Leroy T, Roux C, Villate L, Bodénès C, Romiguier J, Paiva JAP, Dossat C, Aury JM, Plomion C, Kremer A. 2017. Extensive recent secondary contacts between four European white oak species. *New Phytologist* 214:865–878. DOI: 10.1111/nph.14413.
