## Supporting Material File 3 for "Joint analysis of microsatellites and flanking sequences enlightens complex demographic history of interspecific gene flow and vicariance in rear-edge oak populations"

Model A

Summary statistics (see Supporting Material File 2 for definition) listed along the y-axis and value in the x-axis.
Black points: simulated mean value, wide grey line: simulated 95% distribution and fine grey line: 99% simulated distribuiton.
Green, orange or red point: observed suary statistic value that fall within 95% or 99% distribution or fall outside the 99% distriution range, respectively.

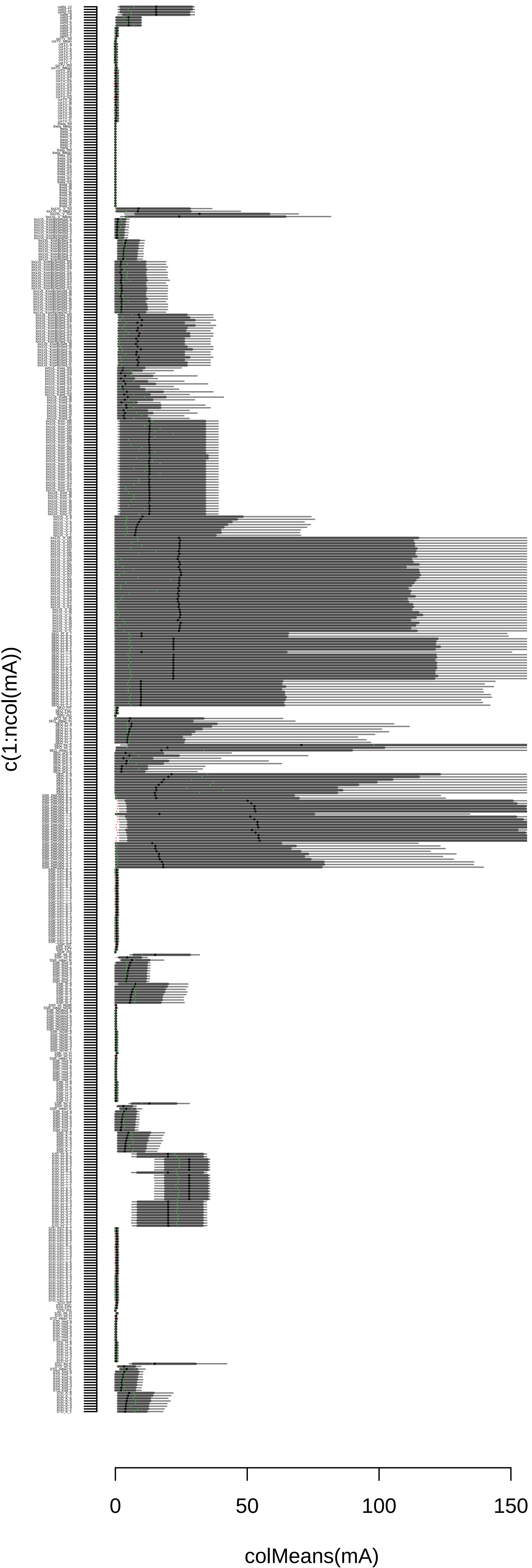

#### Model B

Summary statistics (see Supporting Material File 2 for definition) listed along the y-axis and value in the x-axis.
Black points: simulated mean value, wide grey line: simulated 95% distribution and fine grey line: 99% simulated distribuiton.
Green, orange or red point: observed suary statistic value that fall within 95% or 99% distribution or fall outside the 99% distriution range, respectively.

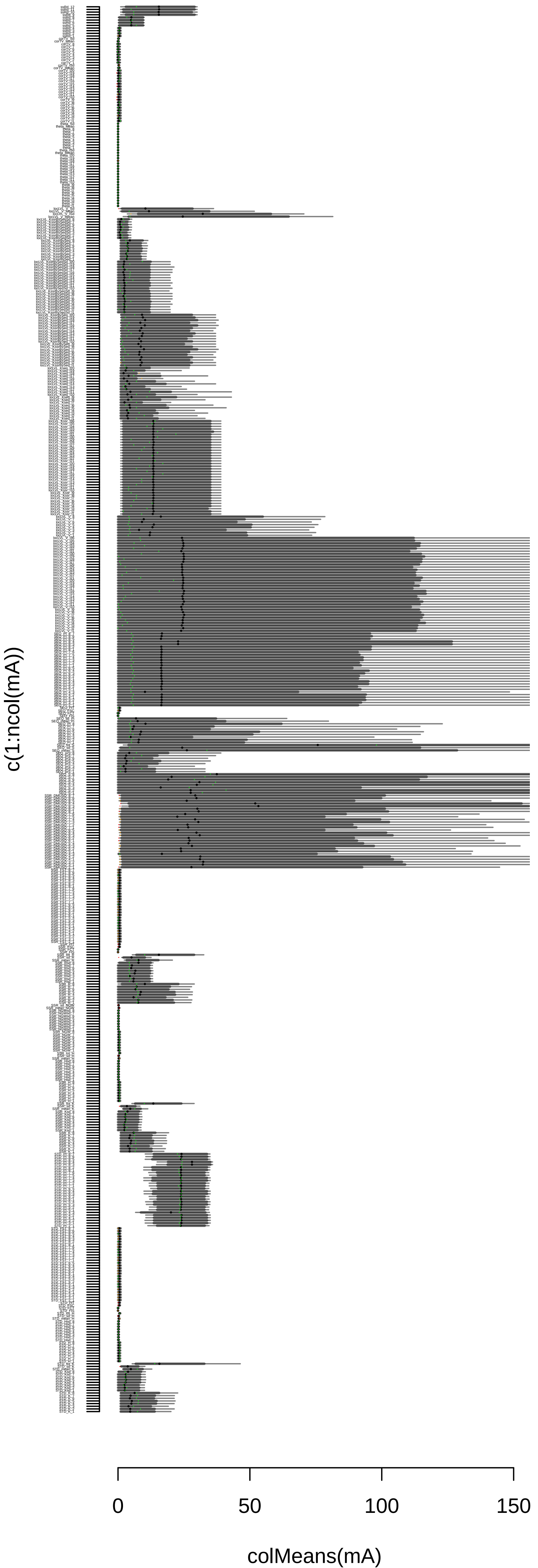

### Model C

Summary statistics (see Supporting Material File 2 for definition) listed along the y-axis and value in the x-axis.
Black points: simulated mean value, wide grey line: simulated 95% distribution and fine grey line: 99% simulated distribuiton.
Green, orange or red point: observed suary statistic value that fall within 95% or 99% distribution or fall outside the 99% distriution range, respectively.

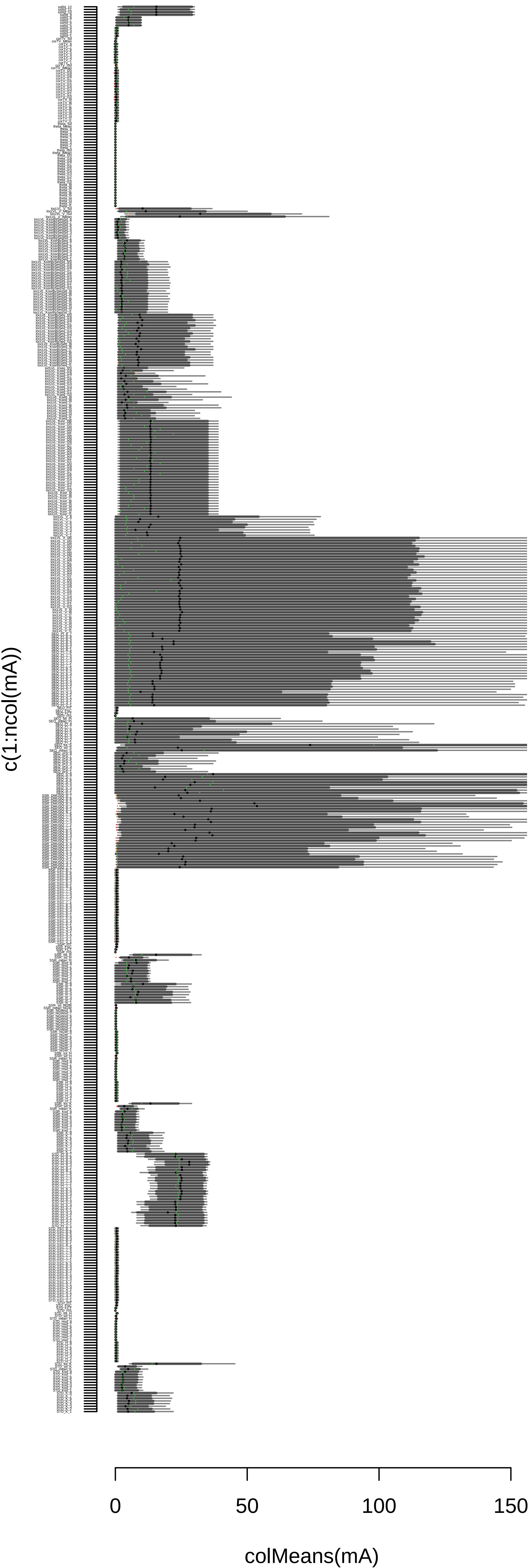

### Model Ab

Summary statistics (see Supporting Material File 2 for definition) listed along the y-axis and value in the x-axis.
Black points: simulated mean value, wide grey line: simulated 95% distribution and fine grey line: 99% simulated distribuiton.
Green, orange or red point: observed suary statistic value that fall within 95% or 99% distribution or fall outside the 99% distriution range, respectively.

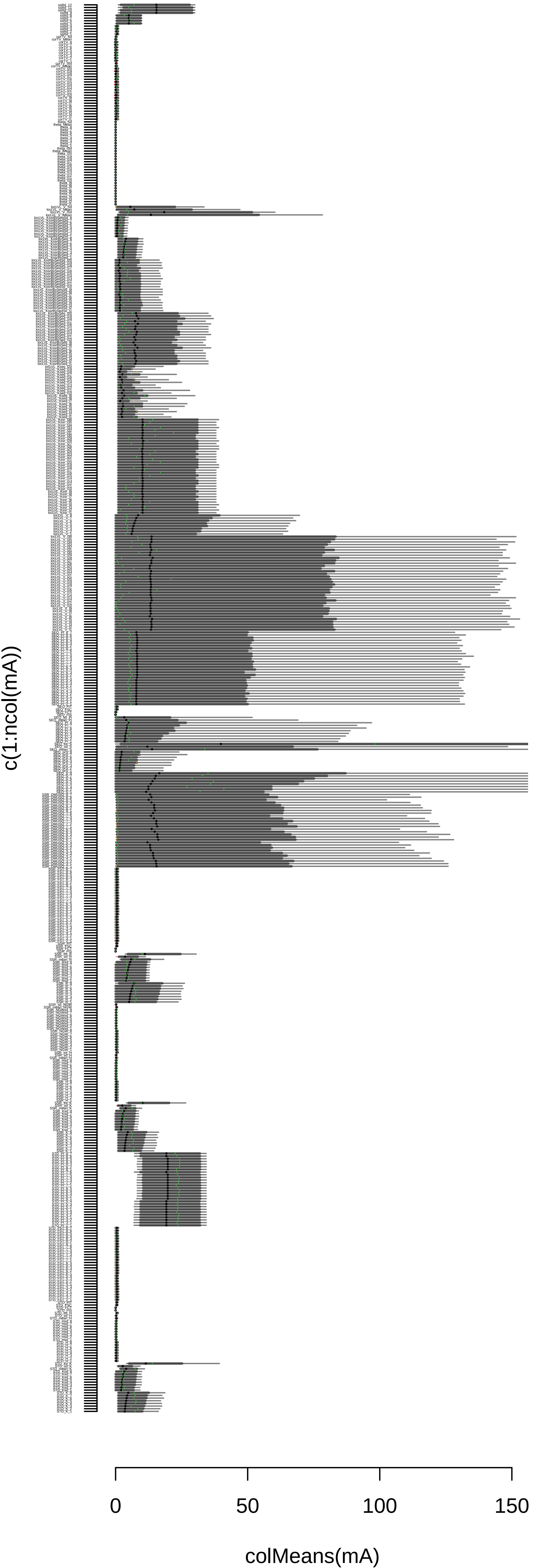

#### Model Bb

Summary statistics (see Supporting Material File 2 for definition) listed along the y-axis and value in the x-axis.
Black points: simulated mean value, wide grey line: simulated 95% distribution and fine grey line: 99% simulated distribuiton.
Green, orange or red point: observed suary statistic value that fall within 95% or 99% distribution or fall outside the 99% distriution range, respectively.

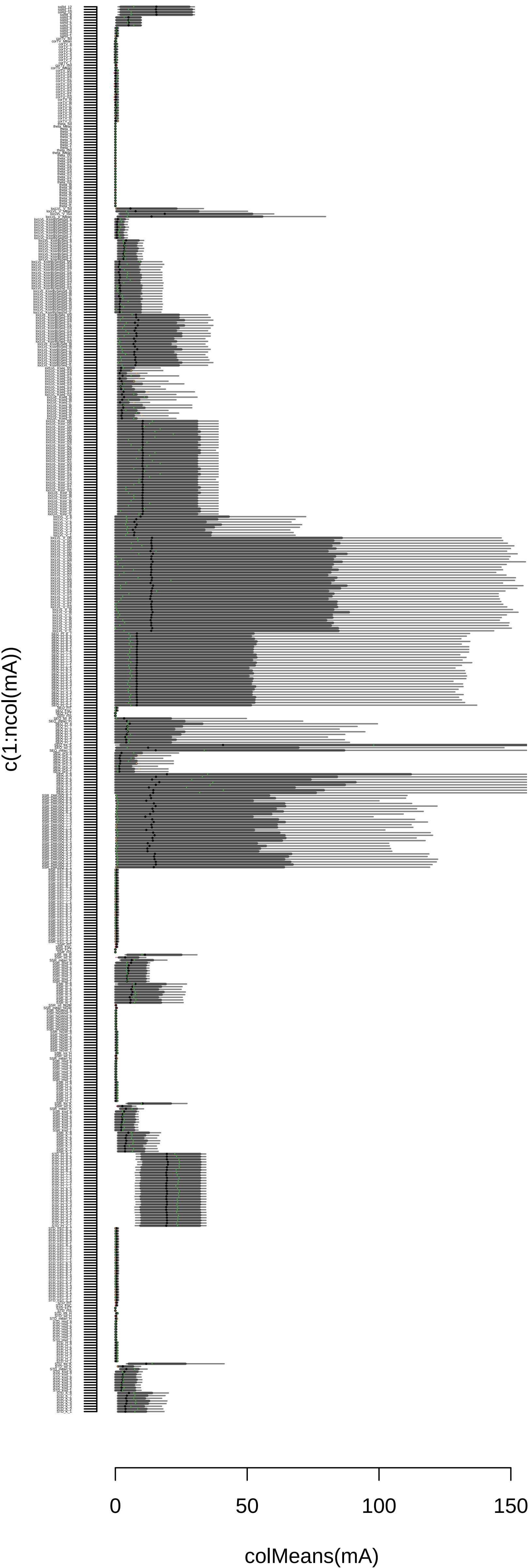

### Model Cb

Summary statistics (see Supporting Material File 2 for definition) listed along the y-axis and value in the x-axis.
Black points: simulated mean value, wide grey line: simulated 95% distribution and fine grey line: 99% simulated distribuiton.
Green, orange or red point: observed suary statistic value that fall within 95% or 99% distribution or fall outside the 99% distriution range, respectively.

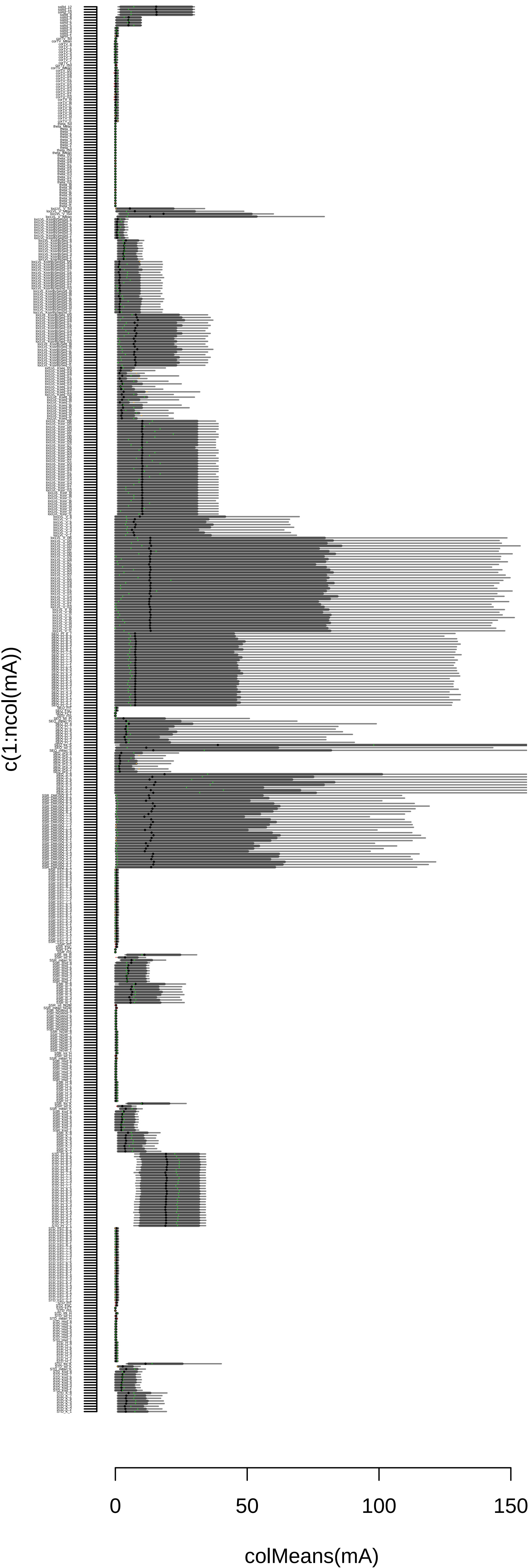
